# Constant sub-second cycling between representations of possible futures in the hippocampus

**DOI:** 10.1101/528976

**Authors:** Kenneth Kay, Jason E. Chung, Marielena Sosa, Jonathan S. Schor, Mattias P. Karlsson, Margaret C. Larkin, Daniel F. Liu, Loren M. Frank

## Abstract

Cognitive faculties such as imagination, planning, and decision-making entail the ability to project into the future. Crucially, animal behavior in natural settings implies that the brain can generate representations of future scenarios not only quickly but also constantly over time, as external events continually unfold. Despite this insight, how the brain accomplishes this remains unknown. Here we report neural activity in the hippocampus encoding two future scenarios (two upcoming maze paths) in constant alternation at 8 Hz: one scenario per 8 Hz cycle (125 ms). We further found that the underlying cycling dynamic generalized across multiple hippocampal representations (location and direction) relevant to future behavior. These findings identify an extremely fast and regular dynamical process capable of representing future possibilities.

## Introduction

Traditional approaches to cognition have focused on the neural representation of external stimuli^1^. In contrast, the ability to generate representations of hypothetical experience, whether of a counterfactual past or a possible future, has only more recently begun to be widely understood as fundamental to the brain^2–8^. This ability, which we here refer to as “generativity,” is essential to a range of cognitive faculties (e.g. planning, imagination, decision-making), indicating a unifying role in cognition. Despite this importance, it remains unclear how generativity is implemented in the brain at the neural level.

Here, behavior and ecology provide direct insight. Generativity contributes to behavior ultimately through projection into the future – future-projection enables outcome prediction, which advantageously guides ongoing behavior. Thus an account of behaviors that entail future-projection can identify biologically necessary properties of generative representation. Crucially, in natural (ecological) settings, high-speed behaviors such as predation and escape are known to require subjects to decide between possible future scenarios not only extremely quickly, but also constantly, as external events continually unfold ^9,10^. This observation implies that the underlying process that generates representations of possible future scenarios has matching properties: sub-second speed and constant operation over time.

Previous work has identified candidate patterns of neural activity encoding possible future scenarios, but these patterns have been found to occur only intermittently and in association with relatively slow (~1 Hz or less), overtly deliberative behaviors, namely head scanning^11,12^ and immobility^13–17^. As a consequence, it has remained unknown how the brain is capable of representing possible future scenarios both quickly and constantly.

## Cycling firing in the hippocampus

To investigate how generativity is implemented in the brain, we sought first to specify a candidate neural substrate. Importantly, generative thinking has been found to activate and require the hippocampus^18,19^, a brain region traditionally linked to memory and spatial navigation. Indeed, recent work on spatially selective hippocampal neurons (place cells) has found activity patterns encoding single generative scenarios in the form of single hypothetical spatial paths^11–17^. Yet despite this advance, these generative activity patterns have been found to occur only intermittently (~1 Hz or slower), and thus cannot implement the speed and constant operation required by natural behavior.

We conjectured that identifying a candidate pattern of neural activity might require a methodological approach inspired by natural behavior: specifically, (i) use of a task that requires generative representation, such as of upcoming spatial paths, and (ii) analysis of neural activity inclusive of periods of high-speed movement. We speculated that these two methods could together prove sufficient if applied to the study of neural substrates implicated in generativity, including the hippocampus.

Taking this approach, we recorded and analyzed hippocampal place cell activity in simple task that requires generative representation: in a bifurcating maze, subjects (rats) allowed to move at high speeds (~50 cm/s) had to choose correctly between two upcoming locations – the left (L) vs. right (R) maze arm – without relying on external directing cues^20–22^ (Fig. 1a, b, **Fig. S1a, b**). Importantly, the bifurcation in the task maze enables unambiguous detection of generative neural activity: in the time period prior to the subject’s choice of arm, neural activity encoding the arm not chosen would necessarily constitute neural activity encoding a possible future scenario (**Fig. S1c, d**). Leveraging this experimental paradigm, we examined the activity of place cells encoding either the L or R arm (**Fig. S1f-h**), focusing on periods of movement prior to subjects’ choice of arm.

**Fig. 1.**
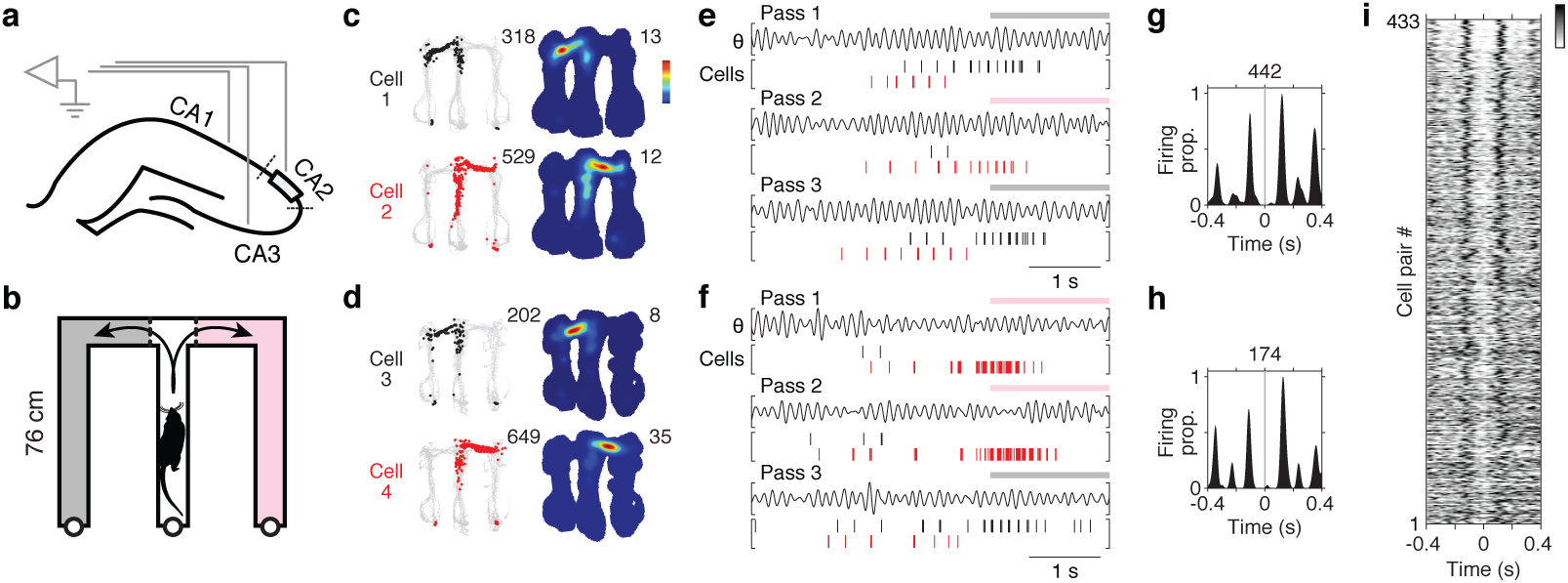
Constant cycling (8 Hz) in the hippocampus. **a**, Diagram of hippocampal recording sites. Recording sites were designated as located in CA2 (grey band) if found to overlap with the CA2 cytoarchitectural locus (see Methods). **b**, Diagram of maze. In the maze task (**Fig. S1b**), subjects (12 rats) were rewarded for correctly choosing between left (L; grey) vs. right (R; pink) maze arms. Arrows illustrate the behavioral choice. Actual maze was not colored differently. **c-d**, Firing maps of two example cell pairs. Each row corresponds to a cell. Data are from outbound maze passes. Left column: positions visited (grey) and positions where the cell fired (colored points; cell 1: black; cell 2: red). Total number of spikes is reported at upper right. Right column: time-averaged firing map. Peak firing rate is reported at upper right. Recording regions by cell: 1: CA3; 2: CA2; 3: CA3; 4: CA3. **e-f**, Firing rasters of the two cell pairs from **c, d** during three maze passes. Plotted above each pass is theta-filtered LFP (θ) (5-11 Hz, CA3). Periods when subject was located in an outer maze arm are indicated by colored patches (grey: left; pink: right). Note the firing alternation between cells at the ~125 ms (8 Hz) timescale. **g-h**, Firing cross-correlogram (XCG) of the two cell pairs from **c, d**. Cell 1 (and 3) spikes are aligned to Cell 2 (and 4) spikes (t = 0 s). Each correlogram (5-ms bins) is smoothed with a Gaussian kernel (σ = 10 ms) and peak-normalized; total number of spikes in XCG is reported at top. **i**, Firing XCGs of anti-synchronous cell pairs (see Methods for criteria). Greyscale value indicates firing density. Data from outbound maze passes; additional cell pair firing types and conditions reported in **Fig. S3**.

Unexpectedly, we found that place cells encoding the L vs. R arms could fire in strikingly regular alternation at ~8 Hz, i.e. a given cell firing on every other 8 Hz cycle (Fig. 1c-h, **Fig. S2a-f**). This pattern of activity was significant in two ways: first, this pattern was unexpected given the classical description of place cells as firing characteristically on adjacent 8 Hz cycles^23–25^ (**Fig. S1n-q**); second, this pattern was the first indication that possible future scenarios (the two upcoming arms) could be encoded with both sub-second speed (i.e. firing within single 8 Hz cycles) and constancy over time (i.e. firing regularly every other 8 Hz cycle).

We further found that place cells encoding opposite heading directions (directional place cells^26^; **Fig. S1i-k**) could fire in regular alternation at 8 Hz (**Fig. S2g-n**), a finding we revisit later. Overall quantification indicated that regular alternation at 8 Hz was a common pattern in the hippocampus (8-9% of cell pair samples, **Fig. S3**). To refer to this dynamical pattern of neural firing, we thereafter used the term “constant cycling,” denoting the combination of regularity and alternation.

Importantly, 8 Hz matches the frequency of hippocampal theta^27,28^, a neural rhythm overtly expressed in the local field potential (LFP) (**Fig. S1l**; shown in Fig. 1, **S2, S3**) and known to entrain hippocampal neural firing^22,29,30^. Indeed theta entrained the firing of the majority of cells in the dataset (90% or 1485 out of 1644 cells; **Fig. S1m-q**), consistent with theta entrainment of 8 Hz cycling and recent work demonstrating theta entrainment of competing neural populations during spatial foraging^31–34^. Given these results, we analyzed periods when theta is continuously active: namely periods of locomotor behavior such as walking and running^29,35^.

## Two correlates of cycling

Prior work has claimed that anatomical^36–38^ and behavioral^29,35,39^ correlates of hippocampal neural activity are essential to understand function. We therefore sought to determine whether cycling firing at 8 Hz had any such correlates. To do so, we analyzed single-cell firing (Fig. 2), for which the 8 Hz cycling dynamic would manifest as “skipped” 8 Hz cycles^33,34^ (cycle skipping) (Fig. 2a-f). Notably, single-cell analysis does not require two particular cells to fire in alternation, thus making fewer assumptions about cell participation from cycle to cycle.

**Fig. 2.**
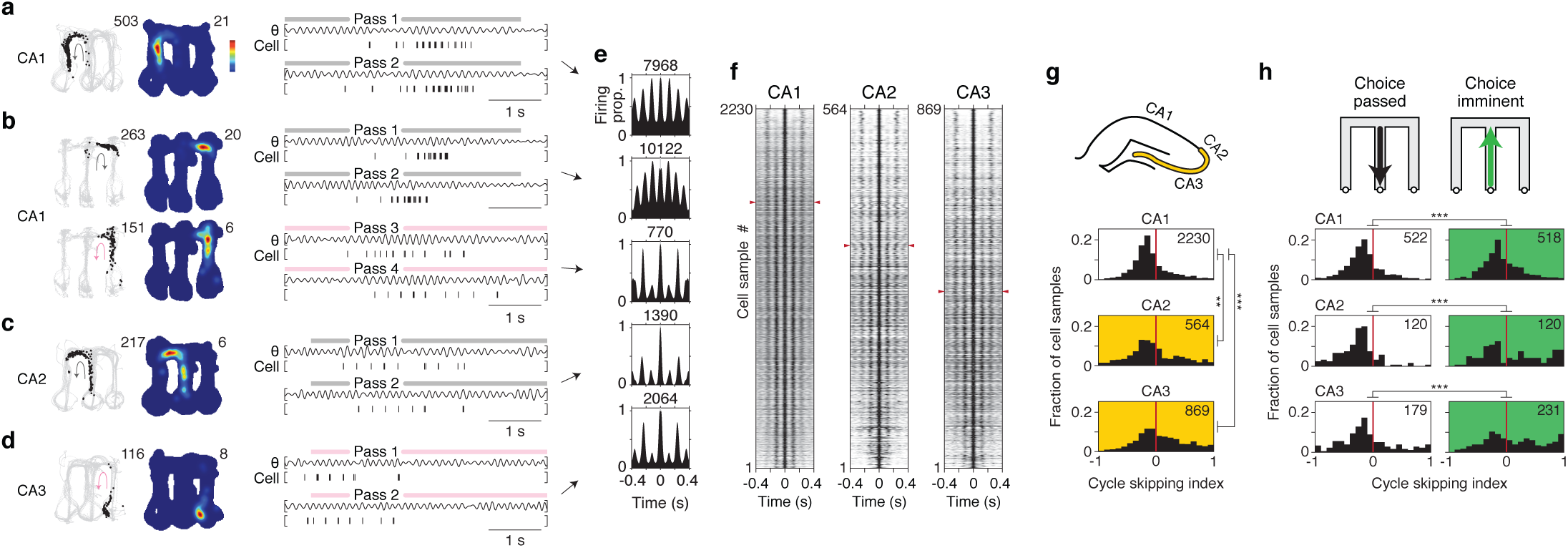
Single-cell correlates of cycling. **a-d**, Time-averaged firing maps (left) and rasters (right) of four example cells (recording regions at far left). Data plotted is exclusively from one type of maze pass (diagramed as arrow; grey: outbound; pink: inbound). In rasters, maze pass times are indicated above plots. Firing maps follow the plotting conventions in Fig. 1c-d. For the cell in **b**, data from two types of maze pass are shown on separate rows to illustrate that firing patterns for an individual cell could depend on condition. **e**, Firing auto-correlograms (ACG) of the four cells from **a-d**. For the cell from **b**, data from each type of maze pass are shown on separate rows. Each ACG (5-ms bins; zero bin excluded) is smoothed with a Gaussian kernel (σ = 10 ms) and peak-normalized. Total number of spikes in ACG is reported at top. **f**, Firing ACGs of all cell samples across recording regions (CA1, CA2, CA3). Greyscale indicates firing density. Each cell sample corresponds to data from a single cell for one type of maze pass. Cell samples are ordered by cycle skipping index (CSI; high to low); for each region, red arrowheads indicate division between cell samples with CSI > 0 (above) vs. < 0 (below). **g**, Cycle skipping index (CSI) by recording region. Top, diagram of sub-regions with higher CSI values (CA2 and CA3; yellow zone). Bottom, CSI histograms. Total number of cell samples is indicated at upper right. Values in CA2/CA3 were higher than in CA1. CA2 vs. CA1, P = 0.0039 CA3 vs. CA1, P = 4.5e-33 **h**, CSI by behavioral condition. Top, schematic of the conditions (choice passed: times when subject was leaving the choice point; choice imminent: times when subject was approaching the choice point). Bottom, CSI histograms. Total number of cell samples is indicated at upper right. For every recording region, values for choice imminent (I) were higher than for choice passed (P). CA1: I vs. P, P = 0.00017 CA2: I vs. P, P = 1.9e-7 CA3: I vs. P, P = 1.1e-5 Wilcoxon rank-sum tests. **, P < 0.01. ***, P < 0.001.

Quantification revealed both an anatomical and a behavioral correlate: (1) cycle skipping was more prevalent in cells recorded in subregions CA2 and CA3 than in CA1 (Fig. 2g, **Fig. S4a, c**); (2) cycle skipping was strongest when subjects approached the choice point, i.e. prior to behaviorally overt choice between maze arms (L vs. R) (Fig. 2h, **Fig. S4b, d)**. This latter finding indicated that behavior-level choice (between upcoming maze paths) recruits the cycling dynamic globally across hippocampal neurons.

## Constant cycling in a neural population

We hypothesized that constant cycling activity, which we first found to encode possible future scenarios in pairs of cells (Fig. 1; **Fig. S2**), might in fact be expressed concurrently across populations of hippocampal neurons. To test this possibility, we analyzed hippocampal neural activity at the population level (Fig. 3). Preliminary inspection of firing across co-recorded cells suggested constant cycling at 8 Hz between possible future locations (L vs. R maze arms) prior to the behavioral choice (**Fig. S5**). For formal analysis, we used a decoding algorithm that is maximally inclusive of the recorded neural firing data (clusterless decoding^40,41^; see Methods). Given the behavior-level correlates identified above (possible future locations in Fig. 1, **Fig. S2a-f** and pre-choice periods in Fig. 2h, **Fig. S4b, d**), we decoded location when subjects approached the choice point. As expected from prior results^42,43^ (**Fig. S1e**), the decoded output showed ~100 ms periods where location projected away from the subject (Fig. 3a-c; **Fig. S6a-f**).

**Fig. 3.**
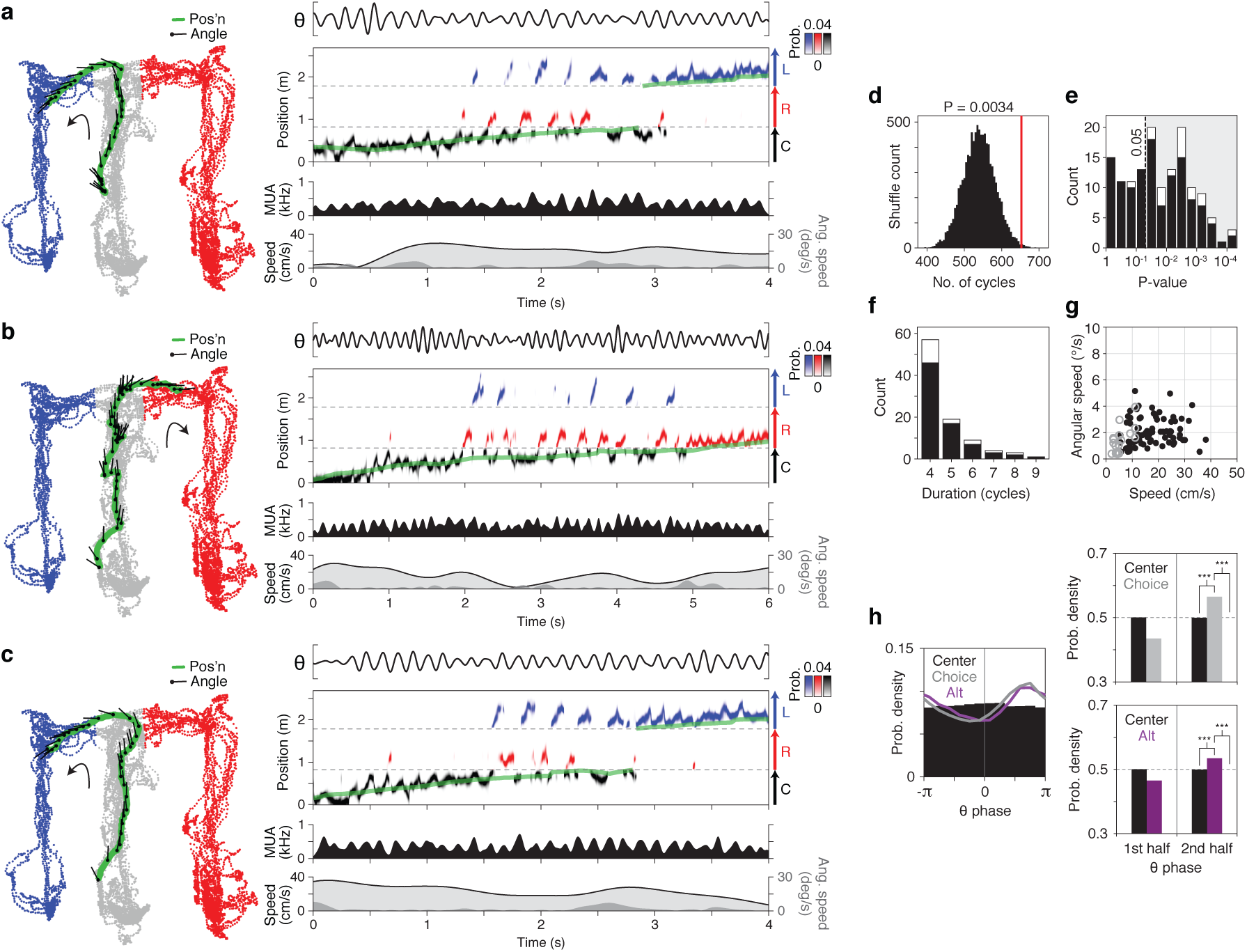
Constant cycling (8 Hz) of possible future locations. Three example maze passes (**a, b, c**) from a single recording epoch. **(Left)** Behavior plot. Position (green) and head angle (black lines; sampling period of plot: 133 ms) are overlaid on positions visited by the subject in epoch (color-coded by maze arm; grey: C (center); blue: L (left); red: R (right)). **(Right)** Data and decoded representation. Top section: LFP (5-11 Hz; CA3). Second section: decoded output (y-axis: linearized position); probability density is plotted as color values and colored by maze arm (black: C; blue: L; red: R); green line indicates actual position of subject. Third section: multi-unit spiking activity (MUA; smoothed with Gaussian kernel (σ = 20 ms)). Bottom section: linear (light grey fill trace) and angular (dark grey fill) speed. **d**, Prevalence of constant cycling in observed (red line) vs. shuffled data (histogram, 10000 permutations). Quantified is the total number of cycles in detected constant cycling periods. P = 0.0038 (38 out of 10000 shuffles had equal or greater prevalence of cycles). **e**, P-values of individual constant cycling periods (temporal shuffle). Shaded area enclosed by dotted line indicates criterion (P < 0.05) for individual periods analyzed subsequently in **f** and **g**; also indicated is whether periods occurred only during movement (>4 cm/s; solid bar) or overlapped with low speed periods (<4 cm/s for <0.5 s; white stacked bar). **f**, Histogram of durations of individual constant cycling periods. Bar plot convention follows that of **e**. **g**, Behavioral speed during individual constant cycling periods. Also indicated is whether periods occurred only during movement (>4 cm/s; solid points) or overlapped with low speed periods (<4 cm/s for <0.5 s; grey circles). Observed periods occurred consistently when angular speed was low (<10°/s), indicating that overtly deliberative behavior^11,12^ was not necessary. **h**, Theta phase distributions of decoded representations (center vs. choice vs. alternative; n = 1683 maze passes from 7 subjects; s.e.m. omitted from plots due to minimal size). Decoded data restricted to center arm. (Left) Mean phase. (Upper right) Choice 2nd half vs. Home 2^nd^ half, P = 4.6e-44, Choice 2^nd^ half vs. 0.5, P = 2.0e-47. (Lower right). Alternative 2nd half vs. Home 2^nd^ half, P = 3.1e-6, Alternative 2^nd^ half vs. 0.5, P = 3.2e-6. Wilcoxon signed-rank tests. ***, P < 0.001.

Remarkably, the decoded output showed periods of constant cycling at 8 Hz between L vs. R maze arms (constant cycling detected as L/R switching across 4 or more successive cycles; Fig. 3a-c, **Fig. S6a-f**), recalling the cycling firing at 8 Hz observed in cell pairs (Fig. 1, **Fig. S2a-f**). To test whether constant cycling between L vs. R representations may have occurred due to random or noisy activity, we shuffled the order of decoded cycles (10000 permutations; cycles segregated with respect to the 8 Hz theta rhythm) and measured the frequency (P-value) with which constant cycling occurred in the shuffled data. This analysis indicated that constant cycling was unlikely to have occurred by chance (P = 0.0034, Fig. 3d; see also **Fig. S6g**). In addition, shuffling within recording epochs identified individual periods of constant cycling that were unlikely to have occurred by chance (93 (of 141 total) constant cycling periods at P < 0.05, Fig. 3e, f; see also **Fig. S6h, i**).

Previous work has described hippocampal neural activity encoding possible future locations as occurring specifically in association with overtly deliberative behavior such as head scans (vicarious trial-and-error^11^); this result led us to ask whether constant cycling at 8 Hz was indeed restricted to such behaviors. We found this not to be the case – constant cycling commonly occurred during high movement speed in the absence of head scans, and thus did not depend on overtly deliberative behavior (Fig. 3g, **Fig. S6j**). We further asked whether decoded activity during movement was reliably associated with upcoming L vs. R choice. We found that the decoded activity did not reliably predict L vs. R (**Fig. S7**), indicating that the cycling dynamic reflected a flexible underlying process not deterministically controlled by overt choice, given the present conditions (navigation/learning prior to asymptotic performance).

## Intra-cycle coding of hypotheticals

How does the hippocampus represent possible futures as fast as at 8 Hz? The finding that 8 Hz cycling was paced by the theta rhythm (Fig. 3, **Fig. S5, 6**) directly suggested that investigating theta would clarify the underlying dynamical process. Specifically, we reasoned that this process had to operate at a timescale finer than full theta cycles: for cycling of future possibilities across theta cycles (inter-cycle coding), there must be a process that generates an individual possibility within each theta cycle (intra-cycle coding).

Intriguingly, classic work^23,44^ has identified an instance of intra-cycle coding: place cells fire at specific phases within theta cycles such that early vs. late phase firing encodes current vs. future location, respectively^23,42–44^. Importantly, this classic case refers only to neural firing related to a single spatial path; this differs from the present study, in which we initially found neural firing related to multiple spatial paths (defined by the maze bifurcation (Fig. 1, **Fig. S2a-f**) or by heading direction (**Fig. S2g-n**)), indicating encoding of hypothetical scenarios. In light of these findings, we conjectured that the classic correlate of intra-cycle coding – current vs. future location – might in fact be an instantiation of a general correlate: namely, current vs. hypothetical scenario. As such, neural firing encoding any particular kind of hypothetical scenario (e.g. locational or directional) would occur at the late phase of theta, akin to firing encoding future location in the classic case^23,44^.

To test this possibility, we first determined whether intra-cycle coding generalizes to representation of possible future locations (L vs. R; **Fig. S1c**) at the population (Fig. 3h) and single-cell (Fig. 4a-d) levels. At the population level, we measured the theta phase when the decoded position represented either the subsequently chosen (choice) or non-chosen (alternative) maze arm. The resulting phase histograms (Fig. 3h, **Fig. S6k**) indicated that the alternative arm was represented selectively at the later phases of theta, equivalently to representation of the choice arm and to future location as in the classic single-path case^42–45^.

**Fig. 4.**
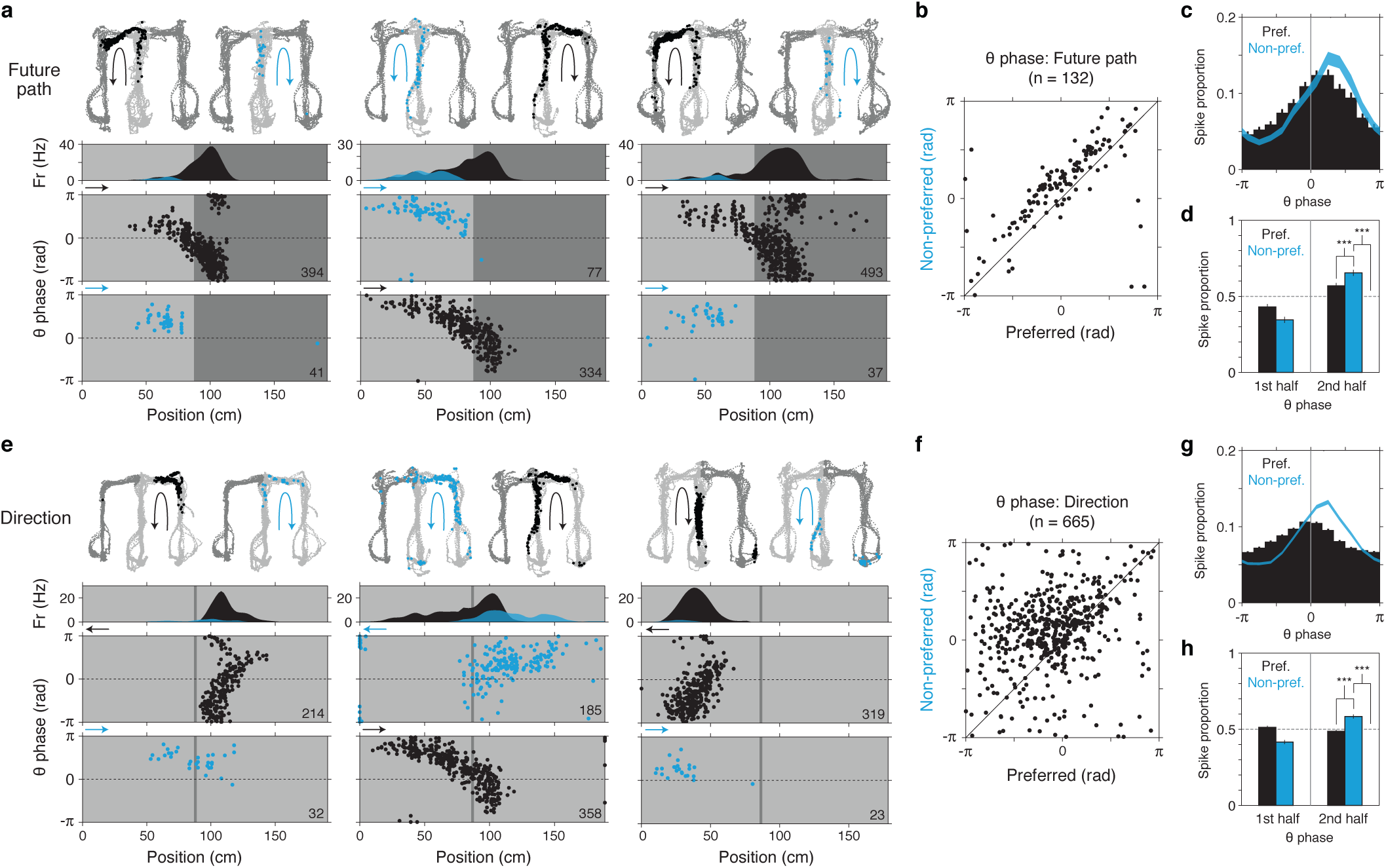
Intra-cycle coding of hypotheticals. Theta phase of firing for future path (**a-d**) and directional (**e-h**) representation in single cells. **a**, Three example cells showing preferred (black) vs. non-preferred (blue) path firing. Each cell is plotted in a column. In each plot, maze locations where subsequent analysis of theta phase of firing (**b-d**) was performed are indicated (light grey; other locations in dark grey). Top section, firing maps. Two maps are shown: one from preferred path (black arrow) maze passes and one from non-preferred path (blue arrow) maze passes. Firing locations during each path pass type are shown as colored points (preferred: black; non-preferred: blue). Second section, time-averaged (linearized) firing map. Bottom section, theta phase of firing by position. Arrow at upper left of plots indicates the subject’s heading direction. Total number of spikes is indicated at lower right. Note that spikes occurring during the non-preferred path tend to occur in the second half of theta (0 to π). **b**, Scatter of mean theta phase. **c**, Theta phase histogram (12-bin). Mean ± s.e.m. **d**, Theta phase histogram (2-bin). Mean ± s.e.m. Note that P vs. NP comparison is expected to depend partially on the location of spatial firing fields. Pref. vs. Non-pref., 2^nd^ half: P = 1.2e-8 Non-pref., 2^nd^ vs. 0.5: P = 2.2e-10 **e**, Three example cells showing preferred (black) vs. non-preferred (blue) directional firing. Plotting conventions follow subpanel **a**. The cell in the middle column is same as that in the middle column in **a**. Note that, as in **a**, spikes occurring during the non-preferred direction tend to occur in the second half of theta (0 to π). **f-h**, Directional firing theta phase quantification. Plotting conventions and comparisons follow **b-d**. Pref. vs. Non-pref., 2^nd^ half: P = 1.6e-35 Non-pref., 2^nd^ vs. 0.5: P = 4.9e-29 Wilcoxon signed-rank tests. ***, P < 0.001.

Importantly, this result yielded further insight into single-cell firing (Fig. 4a-d). Subsets of single place cells are known to fire at different rates depending on which path (e.g. L vs. R) the subject subsequently takes^46–48^, a pattern we here refer to as future-path coding (also known as prospective coding). In light of cycling between differing upcoming locations (Fig. 3), it would be expected that place cells showing future-path coding (higher firing when preferred path is going to be chosen) fire to some extent even when subjects choose the cells’ non-preferred path; furthermore, such non-preferred firing should occur mainly at later theta phases given the population-level result (Fig. 3f). Visual inspection and quantification of single-cell firing confirmed both implications (Fig. 4a-d, **Fig. S8a**). Thus, firing on the non-preferred path is consistent with representation of a possible (hypothetical) future.

Unexpectedly, single-cell analysis indicated that theta phase also governed the well-established hippocampal representation of heading direction^26,49,50^ (schematized and surveyed in **Fig. S1i-k**). Visual inspection (Fig. 4e, **Fig. S8b**) and quantification (Fig. 4f-h) revealed that firing occurring when subjects traveled in cells’ non-preferred direction occurred at later phases of theta, a pattern echoing the future path case (Fig. 4a-d, **Fig. S8a**). In this way, firing in the non-preferred direction is consistent with representation of the non-current, or hypothetical, direction. It is worth noting that past surveys of directional selectivity in single cells have found a markedly wide distribution of selectivity values^50^ (**Fig. S1k**; unlike that of location, **Fig. S1h**), consistent with the possibility that unobserved dynamics govern directional firing.

Further, we also observed equivalent theta phase coding for additional representational firing patterns in the hippocampus (past-path^46–48^ coding and extra-field firing^11^; **Fig. S8c**, **Fig. S9a**). The finding of equivalent temporal organization across multiple neural codes characterized by alternative hypotheticals (of upcoming location (Fig. 3h, Fig. 4a-d) and of direction (Fig. 4e-h)) suggests a single common dynamical process that generates representations of hypothetical scenarios, including possible futures.

## Cycling between directions

The finding that theta phase (intra-cycle) coding – long established for the representation of location^44,45^ – generalizes to non-locational correlates at the single-cell level (Fig. 4, **Fig. S9a**) raises the possibility that theta phase organizes non-locational representations across entire populations of hippocampal neurons.

To determine whether this was the case, we analyzed the hippocampal representation of heading direction, a non-locational variable long known to be a robust correlate of single-cell firing^26,50^ (**Fig. S1i-k**). Initially, we observed instances of population-level activity consistent with intra-cycle organization by inspecting firing in co-recorded place cells grouped by directional selectivity (**Fig. S10a**). For formal analysis, we inferred the representation of direction from population-level hippocampal neural firing using the same decoding approach as applied previously to location (see Methods). Strikingly, the decoded output exhibited periods of <100 ms duration within which the decoded representation switched from current to hypothetical direction and back (**Fig. S9b, c**). Quantification with respect to theta phase revealed that current vs. hypothetical direction were preferentially expressed on the first vs. second halves of theta, respectively, indicating theta phase organization at the population level (**Fig. S9d**). Leveraging this finding, we subsequently decoded in windows that sampled each half of the theta cycle (**Fig. S9b, c**).

This approach revealed periods of constant half-theta cycling of direction (Fig. 5a-c; **Fig. S10b-e**), with comparison to shuffled data (10000 permutations) indicating that constant half-theta cycling was unlikely due to random or noisy activity (P < 0.0001, the lower bound of the test, Fig. 5d). Furthermore, as in constant cycling between possible future locations (Fig. 3h, **Fig. S6k**), periods of constant cycling between current and hypothetical direction (767 (of 12147 total) constant cycling periods at P < 0.05, Fig. 5e-f; **Fig. S10f**) could occur at a wide range of linear and angular speeds (Fig. 5g; **Fig. S10g**), including during running.

**Fig. 5.**
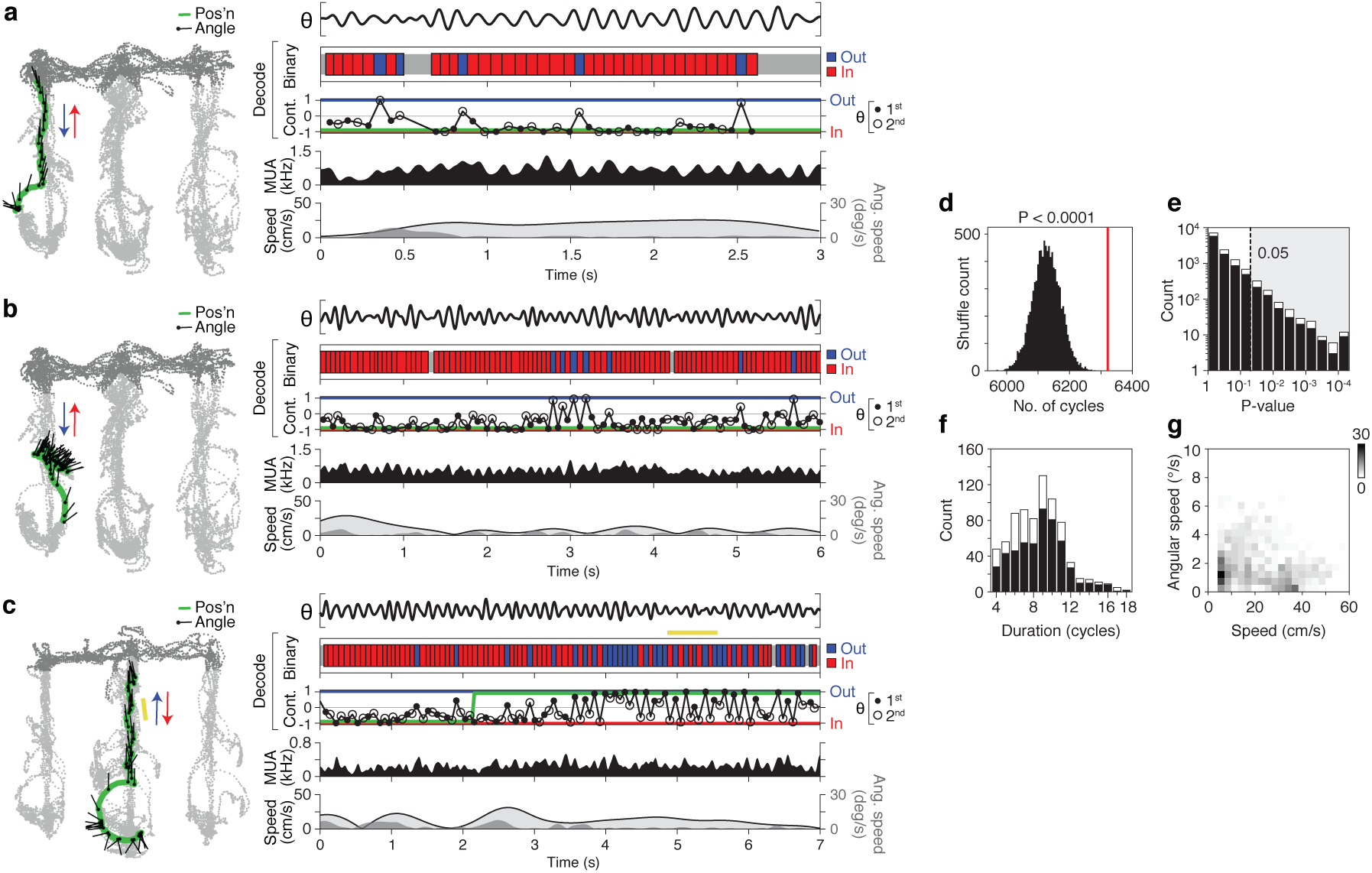
Constant cycling of heading direction. Three example maze passes showing differing amounts of half-theta cycling (**a, b, c**). Examples in **a**, **b** from same recording epoch. In example **c**, an individual period of constant half-theta cycling is highlighted (yellow bar). **(Left)** Behavior plot. Position (green) and head angle (black lines; sampling period of plot: 133 ms) are overlaid on locations visited by the subject in the epoch (light grey: locations analyzed; dark grey: other locations). **(Right)** Data and decoded representation. Top section: LFP (5-11 Hz; CA3). Second section: binary decoded output (red: inbound; blue: outbound). Third section: continuous-valued decoded output (−1: inbound; 0: non-directional; 1: outbound) (filled circle: 1^st^ half theta; open circle: 2^nd^ half theta; connecting lines shown for visual clarity); green line denotes actual direction of subject. Fourth section: multi-unit firing activity (MUA) smoothed with Gaussian kernel (σ = 20 ms). Fifth section: linear (light grey fill trace) and angular (dark grey fill) speed of rat. **d**, Prevalence of constant (half-theta) cycling in observed (red line) vs. shuffled data (histogram, 10000 permutations). Quantified is the total number of cycles participating in detected constant cycling periods. P < 0.0001 (0 out of 10000 shuffles had equal or greater prevalence of cycles). **e**, P-values of individual constant (half-theta) cycling periods (temporal shuffle). Shaded area enclosed by dotted line indicates criterion (P < 0.05) for individual periods analyzed subsequently in **f** and **g**; also indicated is whether periods occurred only during movement (>4 cm/s; solid bar) or overlapped with low speed periods (<4 cm/s for <0.5 s; white stacked bar). **f**, Histogram of durations of individual constant (half-theta) cycling periods. Bar plot convention follows that of **e**. **g**, Behavioral speed during individual constant (half-theta) cycling periods. Greyscale value corresponds to count density (767 total periods plotted). Also indicated is whether periods occurred only during movement (>4 cm/s; solid points) or overlapped with low speed periods (<4 cm/s for <0.5 s; grey circles). Note that constant cycling periods occurred consistently when angular speed was low (<10°/s), indicating that overtly deliberative behavior^12^ was not necessary.

## Discussion

We present two main findings (Fig. 6): (1) the identification of neural activity capable of fast and constant representation of possible future scenarios (constant cycling at 8 Hz) (Fig. 6a); (2) the generalization of the underlying dynamical process across two qualitatively different representations: location and direction (intra-cycle coding of hypotheticals) (Fig. 6b). In addition, we found that cycling dynamics were detectable at three levels of organization (the single-cell (Fig. 2), cell-pair (Fig. 1), and population (Fig. 3) levels), and, further, had both anatomical (Fig. 2g) and behavioral (Fig. 2h) correlates.

**Fig. 6.**
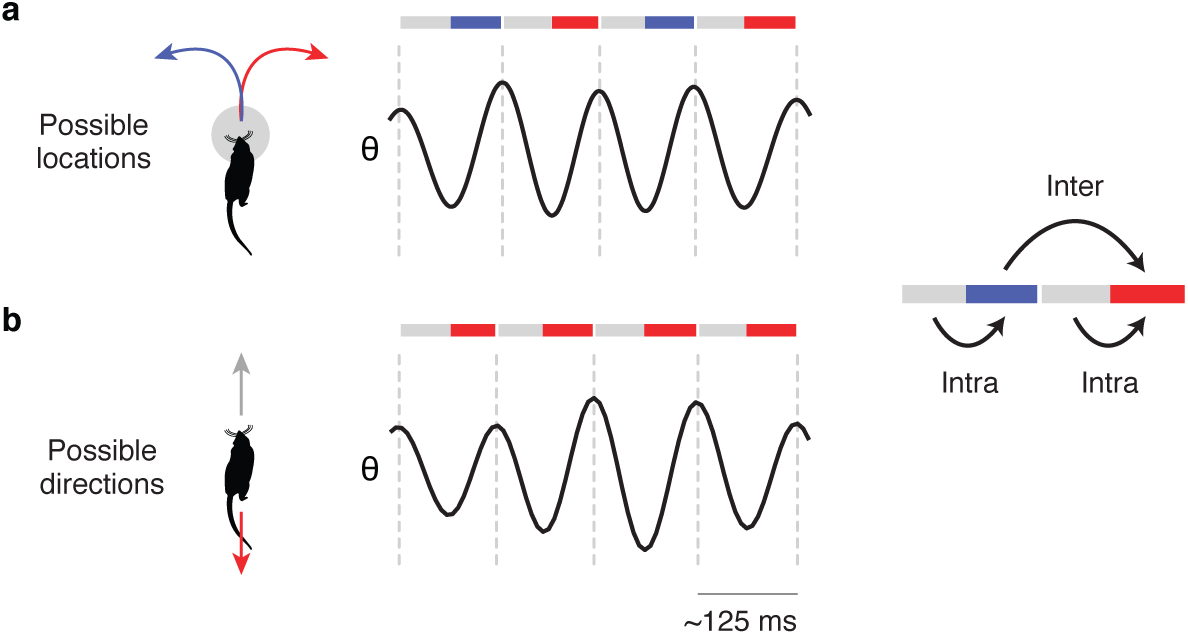
Study summary. Schematic of the two main findings. Each finding refers to one of two patterns of neural firing with respect to the 8 Hz theta rhythm^27–29^ (far right): across successive theta cycles (inter-cycle) or within theta cycles (intra-cycle). The phase of theta is indicated as colored bars (early phase: grey; late phase: red or blue). **a**, Inter-cycle coding of possible future locations (Fig. 1, Fig. 3). Firing at early phases (grey) encodes current location while firing at late phases (red or blue) on successive cycles encodes possible future locations; further, this firing pattern can occur constantly across cycles. **b**, Intra-cycle coding of possible directions (Fig. 4, Fig. 5). Firing at early phases (grey) encodes current direction while firing at late phases (red) encodes possible (hypothetical) direction; further, this firing pattern can occur constantly across cycles. Importantly, this finding generalizes the classic finding referring solely to location (theta phase precession^23,44^), thus indicating that theta phase governs multiple neural codes.

Recent work has established that high-level cognitive functions such as planning and deliberation rely on the brain’s capacity to represent hypothetical (rather than ongoing) experience, a capacity we refer to here as “generativity.” Yet unlike stimulus-driven activity as classically described in sensory neural circuits^1^, the origins of generative neural activity remain poorly understood. Here, place cells in the hippocampus offer a model system given their known generativity – in particular, their representation of hypothetical spatial paths^13,44,51,52^ and spatial contexts^32,53^.

In this study, we find that the rat place cell representation of possible futures has an unexpected conjunction of four properties: segmentation in time (akin to time-division multiplexing in communications systems)^54,55^, sub-second speed (~125 ms and <125 ms), rhythmicity (pacing by the 8 Hz theta rhythm), and generalization of these three dynamical patterns across different representational correlates (location and direction). Identification of these properties specifies putative mechanisms of generativity in three ways: time, neural substrates, and generality.

With respect to time, segmentation within theta cycles (an intra-cycle dynamic^56–59^) specifies a mechanism that can switch between representations of current vs. hypothetical scenarios as fast as double the frequency of theta (equivalent to 16 Hz); in addition, segmentation across theta cycles (an inter-cycle dynamic^31–34^) specifies a mechanism that can switch between representations of hypotheticals as fast as the frequency of theta (8 Hz). The observation of these dynamics builds on previous results demonstrating representational switching at the population level in the hippocampus^11,32,53^, specifically by showing that sub-second rhythmic dynamics governs the representation of behaviorally relevant mutually exclusive possibilities (in the form of alternative future locations and directions), moreover constantly over time. It is also essential to note how these two dynamics accord with an existing framework for current vs. hypothetical representation in the hippocampus^52,60^: the intra-cycle dynamic corresponds to transitions from current (in 1^st^ half-cycle) to hypothetical (in 2^nd^ half-cycle), while the inter-cycle dynamic corresponds to activation of different hypotheticals (across 2^nd^ half-cycles). Furthermore, with regard to external input, both dynamics in the present study occurred in the absence of inducing sensory stimuli (i.e. spontaneously) and under naturalistic conditions (here self-paced, goal-directed navigation); these dynamics were moreover seen in the absence of overtly deliberative behaviors such as directed head scans (vicarious trial-and-error^12^). Most immediately, these various properties suggest revised or new models of network-level theta dynamics^61–66^.

With respect to neural substrates, the rhythmic entrainment of 8 Hz cycling to the theta rhythm implicates brain regions linked to the generation and expression of theta^27,28^. Indeed previous work has reported cycling firing at 8 Hz in subpopulations of neurons in several of these regions (medial septum^67^ and entorhinal cortex^33,34,68^). It is also worth noting that multiple theta generators have been found within the hippocampus itself^5,69^, reminiscent of the finding that cycling firing is expressed at different levels in CA1 vs. CA2/3 neurons (Fig. 2g, **Fig. S4a, c**). These various results suggest privileged roles for structurally defined neural circuits within and beyond the hippocampus. At the same time, the present results implicate a distributed mechanism: cell firing (in single cells or cell pairs) could exhibit cycle skipping in one condition but not another (Fig. 1e, f, Fig. 2b, **Fig. S2, Fig. S3a-d**), indicating that a larger scale of organization is required to explain the generation of cycling.

With respect to generality, the finding that theta dynamics are shared across the representations of location and direction (intra-cycle: Fig. 4; inter-cycle: **Fig. S2**) and likely other representations (**Fig. S9a**), indicates that, at least within the hippocampus, generative activity for different correlates are produced by common theta-associated mechanisms. Indeed, it may be that such theta-associated mechanisms may structure additional (and possibly all) hippocampal representations defined by alternative hypotheticals. Still, it is important to note that the theta rhythm appears to be expressed in a species-specific manner, with a lack of continuous activation during movement in humans, primates, and bats^70^. One intriguing possibility, suggested by recent findings in bats^71^, is that constant cycling could occur without continuous rhythmicity.

Beyond the hippocampus, the present findings imply that neural representation dependent on theta dynamics (intra- and inter-cycle) may be the case in brain regions directly or indirectly connected to the hippocampus^72–74^, including regions associated with mnemonic, evaluative, and executive function; indeed pacing of neural firing by theta has been observed extensively throughout the brain^5,12,27-29,74-77^. More generally, these findings highlight the possibility that neural representation depends on temporal relationships to fast patterns of activity generated within the brain^5,78–80^. Such dependencies may be missed if neural activity is evaluated only over long timescales (e.g. across task trials or over an extended epoch) or only relative to externally observable events (e.g. task cues, behaviors).

The present findings may also have vital implications for the neural basis of decision-making. Previous work suggests that a crucial property of decision-making is speed; in natural environments and with natural behaviors (e.g. predation and escape), decision-making requires a representational mechanism capable of responding to rapidly and continually evolving external conditions (e.g. the behavior of other agents) so that behavior can be re-directed with commensurate speed^9,10^. Sub-second constant cycling dynamics in the hippocampus meet this requirement, raising the possibility that hippocampal activity in particular, and short-timescale neural representation in general, can in fact drive decisions. Consistent with this possibility, single-cell expression of cycling was most prevalent immediately prior to behaviorally reported choice.

Lastly, we note that the present results may also help clarify the neural basis of cognitive functions characterized by generativity^7,8,81^(e.g. recollection, prospection, imagination). Previous work has found that these simulative capacities are associated with activity in a subset of brain regions (including the hippocampus^7,18,19,82-84^), a foundational observation that refers to activity monitored at longer timescales (seconds to minutes). The present findings suggest that (i) methodologically, monitoring activity at the sub-second timescale in these regions may be necessary to resolve competing representations, and (ii) mechanistically, generative representation is coupled to the sub-second dynamics of specific activity patterns: here, the theta rhythm. This mechanistic link may furthermore extend to strikingly similar internally-generated sub-second patterns that govern sensory perception^85–87^ and sensorimotor actions^88,89^; these patterns, currently thought to reflect adaptive mechanisms for sampling information from the external world, may be coordinated with the sub-second patterns of generative activity described here, which can in turn be likened to sampling from internal representations^90,91^.

While the origins, functions, and principles underlying these fast sampling processes remain unclear, formal approaches to cognition highlight the importance of both selectivity and randomness. On the one hand, fully random sampling given finite resources is inefficient and even incompatible with optimal behavior^92^; on the other hand, an overly selective mechanism would prohibit the flexibility, and even the unpredictability^93,94^, suited for dynamic conditions. Future work will need to address whether and how different sub-systems in the brain implement these jointly necessary properties of behaviorally adaptive generativity.

## Acknowledgements

We thank L. Tian, X. Wei, U. Eden, X. Deng, C. Theofanopoulou, A. Palmigiano, and members of the Frank laboratory for comments and discussion, and J. Cunningham and L. Paninski for organizational support during preparation of the manuscript. This work was supported by the Howard Hughes Medical Institute, an NIH grant (R01 MH090188), the NSF NeuroNex Award (DBI-1707398), and the Gatsby Charitable Foundation.

## Author Contributions

K.K. and L.M.F. initially observed the phenomena. K.K., J.E.C., M.S., M.P.K., and M.C.L., collected data. K.K., L.M.F., and D.F.L. designed analyses. K.K. analyzed the data, with J.S.S contributing to preliminary analyses. K.K. and L.M.F. wrote the paper.

The authors declare no competing financial interests.

**Fig. S1. Study basics: localizing vs. generative, maze task, and established findings.**

**a**, A visualization of two types of representation: localizing vs. generative. For illustration, the representational correlate is taken here to be location. Given this correlate, localizing (veridical) representation would refer to current location; in contrast, generative (hypothetical) representation refers to possible locations; for example, sequences of locations (spatial paths) originating from the subject’s current location.

**b**, Schematic of maze task (continuous spatial alternation task^20–22^). The task environment is a W-shaped maze with a center arm and two outer arms. Reward (~0.3 mL of sweetened evaporated milk) is dispensed through 3-cm diameter wells, located at the end of each arm. Rats are rewarded for performing the path sequence shown, in which the correct destination after visiting the center well is the less recently visited outer well.

**c**, Schematic of generative scenarios (possible futures) in the task maze. Diagramed are the two types of outbound maze passes: left and right. When the subject is located in the center arm (white zone) before crossing the choice boundary (dotted lines), entry into each of the two maze arms (choice vs. alternative) constitutes a possible future scenario.

**d**, Identifying generative neural activity: competing hypotheses. For each hypothesis, a schematic of neural activity during a single maze pass is shown; a colored bar indicates the neural activity encoding one of two possible future scenarios: choice (grey) or alternative (blue). The time the subject crosses the choice boundary (CB) is indicated as a dotted line. Neural activity is with reference to a particular brain region of study (e.g. the hippocampus). Note that neural activity can only be generative if the alternative (unchosen option) is encoded (as in iii, iv, v).

i, Pure localizing. Neural activity encodes no information about upcoming experience.

ii, Pure prediction. Before the CB, neural activity only encodes the choice scenario (i.e. fully anticipated experience).

iii, Generative continuous. Before the CB, neural activity encoding the alternative scenario is active continuously.

iv, Generative irregular. Before the CB, neural activity encoding the alternative scenario is active during irregular intervals.

v, Generative regular. Before the CB, neural activity encoding the alternative scenario is active during regular intervals; an underlying rhythmic process is implied.

**e**, Schematic of a fast place cell sequence (encompassing theta and replay sequences^12,42-44,52,95-101^). Diagramed are the spatial firing fields (place fields; colored ovals) of six hippocampal place cells in a single (1-dimensional) path. Fast place cell sequences are ~100 ms population-level firing sequences encoding single spatial paths.

**f**, Schematic of the hippocampal representation of location at a bifurcation. Diagramed are the place fields (colored ovals) of different hippocampal place cells. Under a strictly localizing interpretation^102–104^, place cell activity that occurs before the subject has reached the bifurcation (Center fields: grey) does not encode locations beyond the bifurcation (Left (L) fields: blue; Right (R) fields: red).

**g**, Four example hippocampal place cells showing L vs. R maze arm-specific firing. For each cell, a raw firing map (at left) and a time-averaged firing map (at right) is plotted. Grey points indicate visited locations; black points indicate locations of firing. Total number of spikes (raw map) and peak firing rate (time-averaged map) are shown at upper right.

**h**, Histogram of L vs. R selectivity (all cell samples). Analysis restricted to outbound periods. See Methods for inclusion criteria and definition of selectivity index.

**i**, Schematic of the hippocampal directional representation^26,50,105,106^. Diagramed are the place fields (colored ovals) of different hippocampal place cells that show direction-specific firing (directional place cells). Note that for each heading direction (Outbound (O) or Inbound (I)) one set of cells is predominantly active even though the subject traverses the same locations. For clarity, non-directional cells, which are observed in the same recordings as directional cells, are not shown in the schematic.

**j**, Two example hippocampal cells showing directional firing (O vs. I). For each cell, a raw firing map (upper row; arrow indicates direction) and a time-averaged linearized firing map (bottom row; calculated for maze arm with firing activity) is plotted. For each direction, total number of spikes is plotted at upper right in each raw map. Note that the cell in the first example (at left) exhibits higher direction selectivity than the cell in the second example (at right).

**k**, Histogram of O vs. I selectivity (all cell samples). See Methods for inclusion criteria and definition of selectivity index.

**l**, Example of hippocampal theta (8 Hz) in the local field potential (LFP) during maze. Top row: wide band LFP (0.5-400 Hz; CA3). Middle row: filtered LFP (5-11 Hz). Bottom row: behavioral speed. Note higher amplitude of theta during periods of movement. Far right: power spectral density of LFP during movement (>4 cm/s). Note narrow peak at ~8 Hz corresponding to theta.

**m**, Theta phase of hippocampal cell firing. Firing in single hippocampal cells were histogrammed by theta phase (30° bins, mean ± s.e.m.). Top row, mean phase histogram across cells from each recording region (CA1, CA2, CA3). Cell counts are indicated at top. Bottom row, distribution of modulation depths. The majority of single cells were significantly modulated (90% or 1485 out of 1644 cells; Rayleigh tests at P < 0.05).

**n-q**, Four example cell pairs showing classical 8 Hz firing. Data plotted is from outbound maze path passes (schematized at far left; left (L; grey) vs. right (R; pink) maze arms, with arrows illustrating the behavioral choice). Plotting conventions follow Fig. 1c-e. In these examples, cell firing (in single cells and cell pairs) occurs predominantly on adjacent 8 Hz cycles, as expected in the hippocampus given prior work^24,30,44,45,107^.

**Fig. S2. Cycling firing at 8 Hz: examples.**

Additional example cell pairs showing anti-synchronous cycling firing at 8 Hz.

**a-f**, Six example cell pairs with differing locational representations (left vs. right arm). Plotting conventions are the same as Fig. 1c-g. Data from outbound maze passes (schematic at far left), with left (grey) vs. right (pink) passes plotted separately. **g-l**, Six example cell pairs with differing directional representations. Data from inbound (pink) vs. outbound (grey) maze passes (schematic at far left), with passes of each direction plotted separately. Plotting conventions are the same as Fig. 1c-g but with the following differences: black corresponds to outbound-preferring cells, while red corresponds to inbound-preferring cells; in raw firing maps, colored arrows (black: outbound; red: inbound) indicate corresponding behavior in the maze pass; linearized (rather than 2D) time-averaged firing maps are plotted (arms plotted in linearized maps correspond to positions plotted as light grey in raw firing maps); in rasters, maze pass times are indicated above plots as colored patches. Note that for the example in **l**, two XCGs are plotted, one for each (directional) condition; anti-synchronous cycling firing is seen only in one condition.

**m-n**, Two example cell pairs with similar locational and directional representations.

Data from inbound (pink) vs. outbound (grey) maze passes (schematic at far left, shared with **g-l**), with passes of each direction plotted separately. Plotting conventions are the same as **g-l**, but firing data from cell 26 shown in dark blue to denote outbound preference (same preference as cell 25), and firing data from cell 27 colored in dark red to denote inbound preference (same preference as cell 28). Note that for the example in **n**, two XCGs are plotted, one for each (directional) condition; anti-synchronous cycling firing is seen only in one condition.

**Fig. S3. Cycling firing at 8 Hz: examples and survey of cell pairs.**

**a-d**, Four example cell pairs showing super-synchronous cycling firing at 8 Hz. Plotting conventions follow Fig. 1c-e. Data from outbound maze passes (schematic at far left), with left (grey) vs. right (pink) passes plotted separately.

**e**, Examples of cell pair types (classic and three cycling types) and demonstration of cross-correlogram (XCG) smoothing procedure. Shown are XCGs from four example cell pairs (columnar sections), one of each cell pair type: classic, anti-synchronous, super-synchronous, and off (see Methods for classification criteria). Within each example, columns correspond to time windows of different sizes (left (coarse): ±1 s; right (fine): ± 0.4 s), while rows correspond to raw (upper row) vs. smoothed (lower row; Gaussian kernel: σ = 50 ms (coarse) or 10 ms (fine)) XCGs. Total number of spikes in XCG is reported at top.

**f**, XCGs from outbound maze passes. Greyscale indicates firing density (peak-normalized within each XCG). Coarse and fine XCGs are plotted in paired columns; in addition, all XCGs (at left) vs. cycling-only XCGs (at right) are separately plotted. For both XCG plots (all or cycling-only), XCGs are grouped by cell-pair type (blue: anti-synchronous, yellow: super-synchronous, magenta: off, grey: classic). For the all XCG plot, XCGs are sorted (within cell-pair type) by the timing of the maximum peak in the coarse XCG; for the cycling-only XCG plot, XCGs are sorted (within cell-pair type) by CSI value (highest to lowest).

Total sample counts and percentages (n = 5025): anti: 433 (8.6%; plotted in Fig. 1i), super: 647 (12.9%), off: 306 (6.1%), and regular: 3639 (72.4%). Total sample counts and percentages with exclusion of cell pairs from same tetrode (n = 4251): anti: 381, (9.0%), super: 543 (12.8%), off: 270 (6.4%), and regular: 3057 (71.9%).

**g**, XCGs from inbound maze passes. Same plotting conventions as in **f**.

Total sample counts and percentages (n = 4030): anti: 346 (8.6%), super: 546 (13.5%), off: 211 (5.2%), and regular: 2927 (72.6%). Total sample counts and percentages with exclusion of cell pairs from same tetrode (n = 3409): anti: 290 (8.5%), super: 466 (13.7%), off: 179 (5.3%), and regular: 2474 (72.6%).

**h**, XCGs from four types of maze path passes treated independently (see Methods). Same plotting conventions as in **f**. Total sample counts and percentages (n = 8116): anti: 676 (8.3%), super: 1084 (13.4%), off: 520 (6.4%), and regular: 5836 (71.9%). Total sample counts and percentages with exclusion of cell pairs from same tetrode (n = 6991): anti: 599 (8.6%), super: 928 (13.3%), off, 449 (6.4%), and regular: 5015 (71.7%).

**i**, Histogram of cycle skipping index (CSI) values across cell-pair samples (path-based) and recording regions. Grey shaded regions indicate values corresponding to the classic type (CSI < 0.3); cell-pair samples with CSI > 0.3 were classified as showing cycling firing.

**Fig. S4. Cycling firing at 8 Hz: two correlates.**

Structural correlate (**a**, **c**) and behavioral correlate (**b**, **d**) of cycling firing at the single-cell (**a-b**) and cell-pair (**c-d**) levels. In each panel, results from two criteria for analyzed data are presented (>4 or >20 cm/s locomotor periods; upper and lower sections, respectively); note that cycle skipping (cell CSI > 0; |cell-pair CSI| >~0.3) occurred in either case.

**a**, Single-cell cycle skipping index (cell CSI) by recording region (CA1, CA2, CA3). Same plotting conventions as in Fig. 2g. Two different movement speed thresholds are plotted (>4 cm/s (same as Fig. 2g) and >20 cm/s). Total number of cell samples is indicated at upper right. Values in CA2 and CA3 were higher than in CA1.

CA2 vs. CA1 (>4 cm/s), P = 0.0040

CA3 vs. CA1 (>4 cm/s), P = 4.5e-33

CA2 vs. CA1 (>20 cm/s), P = 0.00032

CA3 vs. CA1 (>20 cm/s), P = 4.4e-21

**b**, Cell CSI by recording region and behavioral condition. Same plotting conventions as in Fig. 2h. Two different movement speed thresholds are plotted (>4 cm/s (same as Fig. 2g) and >20 cm/s). Total number of cell samples is indicated at upper right. Across recording regions, values for choice imminent (I) were higher than for choice passed (P).

CA1: I vs. P (>4 cm/s), P = 0.00017

CA2: I vs. P (>4 cm/s), P = 1.9e-7

CA3: I vs. P (>4 cm/s), P = 1.1e-5

CA1: I vs. P (>20 cm/s), P = 4.1e-7

CA2: I vs. P (>20 cm/s), P = 7.5e-5

CA3: I vs. P (>20 cm/s), P = 0.00061

**c**, Cell-pair CSI (absolute value) by recording region. Same plotting conventions as in **a**. Values in CA2/CA3 were higher than in CA1.

CA2-CA2 vs. CA1-CA1 (>4 cm/s), P = 1.7e-5

CA3-CA3 vs. CA1-CA1 (>4 cm/s), P = 1.2e-9

CA2/3-CA2/3 vs. CA1-CA1 (>4 cm/s), P = 5.9e-19

CA2-CA2 vs. CA1-CA1 (>20 cm/s), P = 0.00017

CA3-CA3 vs. CA1-CA1 (>20 cm/s), P = 0.00029

CA2/3-CA2/3 vs. CA1-CA1 (>20 cm/s), P = 3.3e-12

**d**, Cell-pair CSI (absolute value) by behavioral condition. Same plotting conventions as in **b**. Note that for various regions and in the >20 cm/s condition, the comparison between P vs. I failed to reach significance, possibly due to limiting assumptions underlying the cell-pair (vs. single-cell) CSI; see main text and Methods for discussion.

All-All: I vs. P (>4 cm/s), P = 0.0025

CA1-CA1: I vs. P (>4 cm/s), P = 0.81

CA2-CA2: I vs. P (>4 cm/s), P = 0.38

CA3-CA3: I vs. P (>4 cm/s), P = 0.0018

CA2/3-CA2/3: I vs. P (>4 cm/s), P = 0.00060

All-All: I vs. P (>20 cm/s), P = 0.0045

CA1-CA1: I vs. P (>20 cm/s), P = 0.40

CA2-CA2: I vs. P (>20 cm/s), P = 0.42

CA3-CA3: I vs. P (>20 cm/s), P = 0.61

CA2/3-CA2/3: I vs. P (>20 cm/s), P = 0.039

Wilcoxon rank-sum tests.

*, P < 0.05

**, P < 0.01.

***, P < 0.001.

n.s., not significant (P > 0.05).

**Fig. S5. Constant cycling (8 Hz) at the population level: basic observation.**

Examples of population-level activity indicating constant 8 Hz cycling between representations of possible future locations (left (L) vs. right (R) maze arms). Data from outbound maze passes.

(Far left) Diagram of maze, with L (blue) vs. R (red) maze arms indicated. Actual maze arms were not colored differently. Arrows illustrate the behavioral choice (L vs. R).

(Middle left) Time-averaged linearized firing maps of a population of hippocampal cells (CA1 and CA3 in present example) that were co-recorded. Dotted line indicates location of the choice boundary (CB), beyond which the subject was overtly located in either the Left or Right maze arm. Firing maps are grouped and colored based on the location of peak spatial firing (preferred arm: center, left, or right); in addition, within each arm the cells are sorted by where their peak firing was relative to the center reward well (defined as 0 cm). For each cell, a second firing map corresponding to the non-preferred arm is plotted in the background in a lighter color (of non-preferred arm); slight divergence of the maps at positions before the CB is due to the linearization procedure, which defined the point of divergence between maze arms 10 cm before the CB.

(Right) Data from each of the five maze passes (row sections). Each pass is presented in four column sections (from left to right): (i) full map; (ii) full trace; (iii) highlight trace, and (iv) highlight map.

(i) Full map. Position (green and yellow) and head angle (black lines; sampling period of plot: 133 ms) are overlaid on positions visited by the subject in epoch (colored by maze arm; grey: C (center); blue: L (left); red: R (right)). Highlighted period (yellow) data is expanded in the sections (iii) and (iv).

(ii) Full trace. Top section, theta-filtered LFP (θ) (5-11 Hz, CA3) and times when subject was in an outer maze arm (colored patch; blue: left; right: right). Middle section, firing rasters of cell population. The ordering of the cells follows that of the firing maps (middle left section), with cell groups concatenated. Bottom section, linear (light grey fill trace) and angular (dark grey fill) speed. Highlighted period (yellow) indicated.

(iii) Highlight trace. Time prior to the overt behavioral choice (entry into outer arm) is expanded to show neural activity at the sub-second timescale. Plotting conventions are the same as in (ii), with the addition that times used to segregate a subset of theta cycles in each example are plotted (vertical grey lines in firing raster). Note periods indicating constant cycling (~100 ms / cycle) between Left vs. Right paths in a subset of the 5 example maze passes (from top to bottom, examples 1-3 and 5).

(iv) Highlight map. Plotting conventions are the same as in (i); locations near the CB are expanded to show behavior in greater detail.

**Fig. S6. Constant cycling (8 Hz) between possible future locations: additional examples and approach.**

**a-d**, Examples of constant cycling. Plotting conventions are the same as in Fig. 3a. Panels **a**, **d** are examples of ballistic (uninterrupted) maze passes; **b** is an example of a maze pass interrupted by a stop prior to the choice boundary; **c** is an example of a period of low-speed behavior near the choice boundary.

**e-f**, Examples of constant cycling periods with different decoders: history-dependent vs. uniform prior. Plotting convention is the same as in Fig. 3a, though here with two different decoded outputs: history-dependent (second section) and uniform (third section). Note that either decoder identifies representational cycling, though the two approaches could show differences for individual cycles.

**g-j**, Quantification of decoded output from uniform decoder. Plotting and analysis conventions are the same as in Fig. 3d-g. Study-wide shuffle analysis (panel **g**) indicated that constant cycling was unlikely to have occurred by chance (P < 0.0001, the lower bound of the test; 0 out of 10000 shuffles were equal or greater than the observed prevalence). In addition, shuffle analysis within recording epochs identified individual periods of constant cycling that were unlikely to have occurred by chance (154 (out of 805 total) constant cycling periods at P < 0.05, panels **h-j**).

**k**, Theta phase distributions of decoded representations. Plotting and analysis conventions are the same as in Fig. 3h. (Left) Mean phase.

(Upper right) Choice 2nd half vs. Home 2^nd^ half, P = 1.8e-111, Choice 2^nd^ half vs. 0.5, P = 1.5e-92.

(Lower right) Alternative 2nd half vs. Home 2^nd^ half, P = 2.6e-75, Alternative 2^nd^ half vs. 0.5, P = 2.0e-44.

Wilcoxon signed-rank tests.

***, P < 0.001.

**Fig. S7. Decoding choice.**

Schematic (**a-c**) and results (**d-f**) of three approaches to decoding spatial choice (left (L) vs. right (R) arm). Across approaches, neural activity used to encode was limited to the first half of theta (half-cycle encoder; see Methods), while neural activity used to decode was limited to the second half of theta (half-cycle decoder).

**a**, Approach A (predictive: location encoding). Data used to encode were only from locations beyond the choice boundary (outer maze arms); decoding locations were restricted to locations within the choice boundary (center zone).

**b**, Approach B (predictive: location and path encoding). Data used to encode were from locations throughout the maze; data used to decode were restricted to locations within the choice boundary (center zone). This approach (inclusion of center arm for encoding) allows for localizing neural activity that is path-specific^47,48,108,109^.

**c**, Approach C (non-predictive). Data used to encode and decode were from locations throughout the maze. This approach is shown here as verification that neural activity in present dataset can discriminate between L vs. R spatial choice.

**d-f**, Results from each decoding approach (**a-c**, respectively). Each colored circle corresponds to a single trial (blue: L choice; red: R choice), plotted by actual choice (x-axis, jitter added for visual clarity) and decoded probability of the R choice (y-axis). Also plotted is the mean ± s.d. across trials (filled circle and bar). Reported above plots: (first line) the number of trials for which the decoded choice (R if decoded probability of R was >0.5; L if decoded probability of R was <0.5) matched the actual choice; (second line) P-value of binomial tests (vs. 0.5). For Approach A and B, correspondence between decoded choice (y-axis) and actual choice (x-axis) is equivalent to prediction of upcoming choice in single trials. Note that for either Approach A or B, choice prediction was not reliable (<90%). Overall, it is important to note that the present dataset is of subjects that were introduced to the task and in the process of learning, i.e. prior to asymptotic performance; under highly trained conditions, choice-predictive activity in the hippocampus may be stronger^109,110^.

**Fig. S8. Intra-cycle coding of hypotheticals: additional examples.**

Additional single-cell examples of theta phase organization of preferred vs. non-preferred firing. Examples are grouped by representational correlate (**a**, future path (4 cells); **b**, direction (8 cells); **c**, past path (4 cells)). Plotting conventions are the same as in Fig. 4a, e (spike counts are indicated at lower left in **b**); firing is plotted separately whether occurring in the preferred (black) vs. non-preferred (blue) condition. Note that, across representational correlates, firing in the non-preferred (vs. preferred) condition tends to occur in the second half of theta (0 to π). Firing in the preferred vs. non-preferred condition is consistent with the encoding of current vs. hypothetical experience, respectively (see main text).

**Fig. S9. Intra-cycle coding of hypotheticals: single-cell summary and population decoding.**

**a**, Summary data of theta phase coding for four representational correlates (future path, past path, direction, extra-field; see Methods for definitions). Plotting conventions follow that of Fig. 4b-d, with addition of non-preferred/preferred (NP/P) ratio (third column; ratio of mean non-preferred histogram to mean preferred phase histograms, shown in second column), plotted here to aid comparison. Note that for all representation types, the minimum and maximum theta phase for non-preferred firing are on the first (-π to 0) and second (0 to π) phases of the theta cycle, respectively.

1^st^ row: Future path (n = 132 cell samples)

Pref. vs. Non-pref., 2^nd^ half: P = 1.2e-8

Non-pref, 2^nd^ vs. 0.5: P = 2.2e-10

2^nd^ row: Past path, (n = 21 cell samples)

Pref. vs. Non-pref., 2^nd^ half: P = 0.0037

Non-pref, 2^nd^ vs. 0.5: P = 0.15 (n.s.)

3^rd^ row: Directional (n = 665 cell samples)

Pref. vs. Non-pref., 2^nd^ half: P = 1.6e-35

Non-pref, 2^nd^ vs. 0.5: P = 4.9e-29

4^th^ row: Extra-field (n = 234 cell samples)

Pref. vs. Non-pref., 2^nd^ half: P = 4.7e-22

Non-pref, 2^nd^ vs. 0.5: P = 3.6e-25

Wilcoxon signed-rank tests.

***, P < 0.001.

**b-c**, Examples of directional decoding (showing constant cycling and illustrating decoding windows); each example corresponds to a 1-s segment within the examples in Fig. 4b, c. Each example is divided into the behavioral plot (left section) and time trace (right section). In the behavioral plot, position (green) and head angle (black lines; sampling period of plot: 133 ms) are overlaid on locations visited by the subject in the epoch (light grey: maze locations subject to analysis; dark grey: other locations). In the time trace, six sections are plotted: (i) theta-filtered LFP (5-11 Hz; CA3); (ii) output of sliding-window decoder (−1: inbound; 0: non-directional; 1: outbound); (iii) output of binary decoder (red: inbound; blue: outbound; note that decoding windows are quarter theta cycles centered on each theta half; see Methods) (iv) output of continuous-valued decoder (red: inbound; blue: outbound; filled circle: 1^st^ half theta; open circle: 2^nd^ half theta; connecting lines shown for clarity; actual direction of rat (green line) also shown); (iv) multi-unit firing activity (MUA) smoothed with Gaussian kernel (σ = 20 ms); (v) linear (light grey fill trace) and angular (dark grey fill) speed of rat. Note that the continuous-valued decoder is not equivalent to the mean of the sliding-window decoder, though the decoded outputs are similar: either decoder exhibited constant cycling at the sub-theta cycle timescale (<125 ms).

**d**, Theta phase distribution of decoded directional representations (six plots (1-6) from left to right; data from a single subject in plots 1-3; summary data from seven subjects in plots 4-6). Note that the decoded population-level representation of the hypothetical (non-current) direction was strongest on the second half of theta (0 to pi).

(Plots 1-2) Theta phase histogram of decoded probability density (in (red) vs. out (blue)) when subject was facing in (inbound periods) and out (outbound periods). Decoded posteriors were pooled across all recording epochs.

(Plot 3) Theta phase histogram of decoded probability density (current (black) vs. hypothetical (blue) direction) pooled across all in- and out-bound (facing) periods.

(Plot 4) Theta phase histogram (12-bin, mean ± s.e.m) across subjects (n = 7 rats).

(Plot 5) Hypothetical/current ratio (H/C ratio; ratio of mean current histogram to mean hypothetical phase histogram). Note that the optimal theta phase for current vs. hypothetical decoded density is on the first (-π to 0) vs. second (0 to π) halves of the theta cycle, respectively.

(Plot 6) Mean theta phase histograms (2-bin) for subjects (n = 7 rats). Note that histogram values for each subject are horizontally jittered and offset between groups (current (black) vs. hypothetical (blue)) to show differences between subjects; moreover, within each subject, histogram values (in each theta half) are connected with a line.

Opp. vs. Same, 2^nd^ half, P = 0.016

Opp. 2^nd^ half vs. 0.5, P = 0.016

Wilcoxon signed-rank tests.

*, P < 0.05.

**Fig. S10. Cycling of direction at the population level.**

**a**, Example population-level firing indicating cycling dynamics in the representation of heading direction (outbound (out) vs. inbound (in)).

**(Far left)** Diagram of maze and alternative directions (out: black; in: blue).

**(Middle left**) Time-averaged linearized firing maps of a co-recorded population of 20 hippocampal cells (CA1, CA2, CA3). Firing maps are grouped and colored based whether their peak firing was in the out (Out, black) vs. in (In, blue) condition, with non-directional cells (Non, black) plotted with the out group; in addition, within each group the cells are sorted by the position of their peak firing relative to the center reward well (0 cm). For each cell, a second firing map corresponding to the non-preferred direction is plotted in background in a lighter color. In this example, cells 1-13 either preferred to fire in the outbound direction (cells 4, 5, 7-9, 12) or fired similarly in either direction; cells 14-20 preferred to fire in the inbound direction.

(**Right**) Example data from three maze passes (row sections). For each maze pass, data is presented in four column sections (from left to right): (i) full map; (ii) full trace; (iii) highlight traces.

(i) Full map. Position (green and yellow) and head angle (black lines; sampling period of plot: 133 ms) are overlaid on positions visited by the subject in epoch. Highlighted period (yellow) data is expanded in the following sections.

(ii) Full trace. Top section, sign-reversed theta-filtered LFP (-θ) (5-11 Hz, CA3; sign inversion to approximate theta as measured in CA1^69^). Middle section, firing rasters of cell population. The ordering of the cells follows that of the time-averaged firing maps, with cell groups concatenated. Bottom section, linear (light grey fill trace) and angular (dark grey fill) speed. Highlighted periods (yellow) indicated.

(iii) Highlight traces. Plotting conventions are the same as in (ii). Note instances of activation of multiple inbound-preferring cells, indicating population-level representation of the inbound (hypothetical) direction; further note that these activations occur on the second half (rising phase; 0 to pi) of the theta-filtered LFP.

b-e, Examples of constant (half-theta) cycling of direction. Plotting conventions are the same as in Fig. 5a.

**f**, **g**, Quantification of constant (half-theta) cycling of direction with minimum of 8 (half-theta) cycle criterion. Analysis procedure is the same as Fig. 5e-g (individual constant cycling period shuffle; see Methods), with additional requirement of a minimum of 8 half-theta cycles for detected constant cycling periods; 483 (of 820 total) constant cycling periods were detected. Plotting conventions in **f** and **g** are the same as that in Fig. 5e, g, respectively.

## Methods summary

Single neuron (cell) activity and local field potentials (LFP) were recorded from CA1, CA2, CA3, and DG in the dorsal hippocampus of twelve male rats performing a continuous spatial alternation task^20–22^.

## Methods

### Subjects, neural recordings, and behavioral task

Twelve male Long-Evans rats that were 4 to 9 months old (500–600 g) were food deprived to 85% of their baseline weight and pre-trained to run on a 1-m linear track for liquid reward (sweetened evaporated milk). After subjects alternated reliably, they were implanted with microdrives containing 14 (two subjects), 21 (three subjects), 25 (one subject), or 30 (six subjects) independently movable four-wire electrodes (tetrodes^104,111^) targeting dorsal hippocampus (all subjects) and medial entorhinal cortex (two subjects), in accordance with University of California San Francisco Institutional Animal Care and Use Committee and US National Institutes of Health guidelines. Data from all subjects have been reported in earlier studies^21,22,112,113^.

In five subjects, right and left dorsal hippocampus were targeted at AP: −3.7 mm, ML: ± 3.7 mm. In two subjects, dorsal hippocampus was targeted at AP: −3.6 mm, ML: +2.2 mm, in addition to medial entorhinal cortex at AP: −9.1, ML: +5.6, at a 10 degree angle in the sagittal plane. In five subjects, right dorsal hippocampus was targeted at AP: −3.3 to −4.0 mm, ML: +3.5 to +3.9 mm, moreover, in two of these subjects, the septal pole of right hippocampus was targeted with an additional six tetrodes targeted to AP: −2.3 mm, ML: +1.1 mm. Targeting locations were used to position stainless steel cannulae containing 6, 14, 15, or 21 independently driveable tetrodes. The cannulae were circular except in four cases targeting dorsal hippocampus in which they were elongated into ovals (major axis ∼2.5 mm, minor axis ∼1.5 mm; two subjects with major axis 45° relative to midline, along the transverse axis of dorsal hippocampus; two subjects with major axis 135° relative to midline, along the longitudinal axis of dorsal hippocampus). Data from tetrodes targeting both right and left dorsal hippocampus were analysed in this study.

In five subjects, viral vectors with optogenetic transgenes were targeted to either right dorsal CA2 (three subjects, AAV2/5-CaMKII-hChR2(H134R)-EYFP, UNC Vector Core, 135 nl at AP: −3.6 mm, ML: +4.2 mm, DV: −4.5 mm), dorsal DG (one subject, AAV2/5-I12B-ChR2-GFP (see ref. 52 for details about the I12B promoter)), 225 nl at AP: −3.75 mm, ML: +2.2 mm, DV: 3.9 mm and AP: −3.75 mm, ML: +1.8 mm, DV: −4.5 mm), or right supramammilary nucleus (one subject, AAV2/5-hSyn-ChETA-EYFP, Penn Vector Core, 135 nl at AP: −4.3 mm, ML: +1.8 mm, and −8.9 mm along a trajectory angled at 6° in the coronal plane). Viruses were delivered during the implant surgery using a glass micropipette (tip manually cut to ∼25 µm diameter) attached to an injector (Nanoject, Drummond Scientific). In addition, a driveable optical fiber (62.5/125 µm core/ cladding) was integrated in the tetrode microdrive assembly to enable light delivery to hippocampus. This fibre was advanced to its final depth (2.5–3 mm) within 7 days of implantation. Data reported in this study were collected before light stimulation. No overt differences in neural activity were observed in subjects that received virus^22^.

Over the course of two weeks following implantation, the tetrodes were advanced to the principal cell layers of CA1 (all subjects), CA2 (5 subjects), CA3 (11 subjects), and DG (3 subjects). For DG, tetrodes were advanced to the cell layer using a previously described protocol in which the tetrodes were slowly advanced within DG (~10 µm increments) and unit activity monitored over long periods of rest^114^. DG cell layer was identified by the presence of highly sparsely firing putative principal units. In several subjects, tetrodes were also left in cortex overlying dorsal hippocampus. Neural signals were recorded relative to a reference tetrode positioned in corpus callosum above right dorsal hippocampus. The reference tetrode reported voltage relative to a ground screw installed in skull overlying cerebellum, and local field potential (LFP) from this tetrode was also recorded. All tetrode final locations were histologically verified (see below).

After 5–7 days of recovery after surgery, subjects were once again food deprived to 85% of their baseline weight, and again pre-trained to run on a linear track for liquid reward. At ~14 days after surgery, six subjects were then introduced to one task W-maze and recorded for 3 to 6 days before being introduced to a second task W-maze, located in a separate part of the recording room and rotated 90° relative to the first. On recording days in which the second task W-maze was used, recordings were also conducted in the first task W-maze. In two subjects, recordings were conducted in both task W-mazes on every recording day. The W-mazes were 76 × 76 cm with 7-cm-wide track sections. The two task W-mazes were separated by an opaque barrier.

In each W-maze, subjects were rewarded for performing a hippocampus-dependent continuous alternation task (**Fig. S1b**). Liquid reward (sweetened evaporated milk) was dispensed via plastic tubing connected to a hole at the bottom of each of the three reward wells, miniature bowls 3 cm in diameter. In eight subjects, reward was dispensed via syringes operated manually by an experimenter who was located in a separate part of the recording room. In five subjects, entry of the subject’s head into reward wells was sensed by an infrared beam break circuit attached to the well, and reward was automatically delivered by syringe pumps (OEM syringe pumps, Braintree Scientific) either immediately or after an imposed delay lasting from 0.5 to 2 s. Task epochs lasting 15 min were preceded and followed by rest epochs lasting ~20 min in a high-walled black box (floor edges 25–35 cm and height 50 cm), during which rats often groomed, quietly waited, and slept. Two subjects also ran in an open field environment for scattered food (grated cheese) after W-maze recordings, with additional interleaved rest epochs. Tetrode positions were adjusted after each day’s recordings.

Data were collected using the NSpike data acquisition system (L.M.F. and J. MacArthur, Harvard Instrumentation Design Laboratory). During recording, an infrared diode array with a large and a small cluster of diodes was affixed to headstage preamplifiers to enable tracking of head position and head direction. Following recording, position and direction were reconstructed using a semi-automated analysis of digital video (30 Hz) of the experiment. Spike data were recorded relative to the REF tetrode, sampled at 30 kHz, digitally filtered between 600 Hz and 6 kHz (2-pole Bessel for high- and low-pass), and threshold crossing events were saved to disk. Local field potentials (LFPs) were sampled at 1.5 kHz and digitally filtered between 0.5 Hz and 400 Hz. LFPs analysed were relative to the REF tetrode except where otherwise indicated.

Single-cell (unit) neural firing was identified by clustering spikes using peak amplitude, principal components, and spike width as variables (MatClust, M.P.K.). Only well-isolated units with stable spike waveform amplitudes were clustered. A single set of cluster bounds defined in amplitude and width space could often isolate units across an entire recording session. In cases where there was a shift in amplitudes across time, units were clustered only when that shift was coherent across multiple clusters and when plots of amplitude versus time showed a smooth shift. No units were clustered in which part of the cluster was cut off at spike threshold.

### Histology and recording site assignment

After recordings, subjects were anesthetized with isoflurane, electrolytically lesioned at each tetrode (30 µA of positive current for 3 s applied to two channels of each tetrode), and allowed to recover overnight. In one subject, no electrolytic lesions were made, and tetrode tracks rather than lesions were used to identify recording sites. Subjects were euthanized with pentobarbital and were perfused intracardially with PBS followed by 4% paraformaldehyde in PBS. The brain was post-fixed *in situ* overnight, after which the tetrodes were retracted and the brain removed, cryoprotected (30% sucrose in PBS), and embedded in OCT compound. Coronal (10 subjects) and sagittal (2 subjects) sections (50 µm) were taken with a cryostat. Sections were either Nissl-stained with cresyl violet or stained with the fluorescent Nissl reagent NeuroTrace Blue (1:200) (Life Technologies, N-21479). In four subjects, the sections were blocked (5% donkey serum in 0.3% Triton-X in TBS, used for all incubations) for 1 h, incubated with RGS14^115–117^ antibody (1:400) (Antibodies Inc., 75-140) overnight, washed, and subsequently incubated with fluorescent secondary antibody (1:400) (Alexa 568, Life Technologies). CA2 recording sites were designated as those in which the electrolytic lesion or end of tetrode track overlapped with the dispersed cytoarchitectural zone characteristic of CA2^73,116,118-121^. It is important to note that CA2 sites defined in this way include recording locations that have been designated in previous studies as ‘CA3a’.

### Data analysis

All analyses were carried out using custom software written in Matlab (Mathworks).

### Cell inclusion and classification

Units (single cells) analyzed in the study were those that fired at least 100 spikes in at least one task epoch and had at least 50 spikes in their auto-correlogram (0-40 ms; t = 0 excluded). Across all cells, a scatter plot of mean firing rate (from task epoch with highest mean rate), spike width, and autocorrelation function mean (0-40 ms; low values indicating burst firing) showed two clusters^22,30,122-126^. Putative principal cells corresponded with the low firing rate (<4 Hz), large spike width, low autocorrelation mean cluster, while putative interneurons corresponded to the cluster characterized by high firing rate, small spike width, and high autocorrelation mean. Thirty-seven cells with ambiguous features were left unclassified.

Total putative principal unit counts across recording sites were CA1: 978, CA2: 250, CA3: 528, DG: 17. Following previous work^22,127^, a subset of CA2-site putative principal cells were identified by whether they were positively modulated (CA2 P cells) vs. non-positively modulated (CA2 N cells) by the sharp wave-ripple network pattern; as firing activity in both cell types overlaps with theta-modulated (locomotor) periods^22^, both were included in analysis.

### Maze linearization (segments, arms, choice boundary, center zone)

For later analyses, the 2D position of subject was converted into 1D position (linearized position) along the three arms (center, left, and right arms) of the task maze. The three arms meet at the central junction, with the center arm composed of a one linear segment, and the left and right arms composed of two linear segments. Linearization requires specification of the six segment endpoints (2D coordinates; three corresponding to the locations of the three reward wells, and three corresponding to the junctions between linear segments); these endpoints were specified manually prior to analysis of neural data.

Linearized position was obtained by projecting the 2D position of subject onto the nearest linear segment and measuring the distance, along the maze arms, from the center well (defined as 0 cm). Note that every positional data point is also assigned one of five maze segments and one of three maze arms.

The choice boundary was defined as the linearized position 10 cm beyond the central junction; the choice boundary was interpreted as the position where spatial choice (left vs. right) for outbound maze paths (**Fig. S1b**) was reported behaviorally (overt). The center zone was defined as the set of linear positions (mainly correspondent with the center arm) prior to choice boundary. Note that the choice boundary refers to the boundary with either the L or R arm (e.g. either of the two interfaces between color-coded regions in behavioral maps of Fig. 3).

### Behavioral states: movement, maze, and task

Analyses in this study refer to periods of time defined by subjects’ behavior, whether with respect to movement, the spatial layout of the environment (maze), or to the structure of the continuous alternation task.

#### Locomotion

Locomotor periods were defined as times when head speed was >4 cm/s. Single-cell and cell-pair analyses were restricted to locomotor periods, while decoding analyses included additional 0.5 s flanking periods, described further below.

#### Task paths

Task paths were defined as the four spatial trajectories relevant to the task (two outbound: Center to Left and Center to Right; two inbound: Left to Center and Right to Center; **Fig. S1b**).

#### Direction

Locomotor periods were fully subdivided into two types of periods defined by heading direction: outbound or inbound. Outbound vs. inbound periods were times when the subject’s head direction was aligned to outbound vs. inbound task paths (given the assigned maze segment; see above), respectively; furthermore, alignment was binary: the assigned direction (outbound vs. inbound) was the one yielding the smaller angle (0 to 180°) along the maze segment.

#### Maze passes and path passes

Maze passes were defined as single contiguous periods where the animal traveled from one reward well to another (or returned to the same well); these periods were comprised largely of locomotor periods but also included occasional intervening non-locomotor periods (i.e. head scans^12,128^ and stops). Maze passes were classified into nine types: four corresponding to the four task paths (path passes; two outbound and two inbound), two corresponding to traversals between the two outer wells (left to right and right to left), and three corresponding to backtracking traversals in which the subject left one of the three wells and returned to that same well without reaching a different well. Backtrack passes were detected only if the pass lasted at least 2 s and the subject traveled at least 20 cm from the well in linearized position. The start and end of maze passes were times when the linear position of the subject diverged from and met the linearized positions of the start and end wells, respectively.

#### Path periods

Path periods were defined as the subset of times within path passes when (i) the subject was located in one of the three maze segments defining the current task path and (ii) the subject’s heading direction was the same as (aligned to) the direction defining the current task path.

### Spatial firing in single cells

Spatial firing was quantified using linearized (1D) position, with plots of 2D spatial firing maps (e.g. Fig. 1) only for illustration. If data from two unique maze environments were available for a cell, and the cell fired at least 100 spikes in the both environments, then the cell was analyzed in the two unique maze environments independently. Thus cells in the dataset could contribute more than one sample to subsequent analyses (cell samples). If multiple recording epochs in one maze environment were available, then the cell was analyzed in recording epoch for which the cell had the highest mean firing rate.

Following the established direction^26,50,106^ and path^46-48,109^ dependence of hippocampal spatial firing, analysis was performed on locomotor data subdivided in two ways, either (I) direction or (II) path. For (I), analysis was performed separately for outbound vs. inbound periods, moreover separately for each of the three maze arms; thus a given cell in a given maze environment could contribute up to three samples in subsequent direction-based analyses (direction-based cell samples; outbound and inbound). For path, analysis was performed separately for periods corresponding to the four task path types (for which a minimum of five available passes were required for each path type); thus a given cell in a given maze environment could contribute up to four samples in subsequent path-based analyses (path-based cell samples; outbound left, outbound right, inbound left, and inbound right). Two days of recordings for one subject were excluded from analysis for lack of path passes in both outer maze arms.

For each cell sample, a time-averaged firing map was calculated for each period type. First, total spike counts and occupancy durations were tabulated in 2-cm bins. Both occupancy and spike counts per bin were smoothed with a Gaussian window (σ = 4 cm), then spike counts were divided by occupancy to produce the cell’s smoothed occupancy-normalized firing map. In sporadic cases, spatial bins with insufficient occupancy (<50 ms in a 2-cm bin) were excluded from analysis. For each time-averaged firing map, the peak (spatial) firing rate was defined as the maximum value across position bins; spatial firing fields (place fields) were detected as sets of contiguous positions with rate >2 Hz and at least 10 cm large.

Note that in the case of direction-based (I) firing maps (e.g. Fig. 4e, **Fig. S2g-n**), the spatial bin at the center junction appears as a minimum as a result of the above procedure of calculating firing maps for each of the three maze arms separately. For plots, the value of this spatial bin was linearly interpolated; this value was otherwise excluded from subsequent quantification.

Two-dimensional time-averaged firing maps (plotted for illustration) were calculated with an analogous procedure using 1-cm square bins and a symmetric 2D Gaussian smoothing window (σ = 3 cm).

### Selectivity index

To measure the specificity of representation at the single-cell level, a selectivity index was calculated, specifically one for location (Left (L) vs. Right (R) arms; survey in **Fig. S1h**) and one for heading direction (Outbound (O) vs. Inbound (I); survey in **Fig. S1k**).

If data from two unique maze environments were available for a cell, and the cell fired at least 100 spikes in both environments, then the cell was analyzed (selectivity indices calculated) in the two unique maze environments independently. Thus cells in the dataset could contribute more than one cell sample to subsequent analyses (cell samples). If multiple recording epochs in one maze environment were available, then the cell was analyzed in recording epoch for which the cell had the highest mean firing rate.

The location selectivity index was calculated for cell samples for which the 1D firing maps that had at least one place field in either the L or R maze arm. To assess cell activity directly relevant to the spatial choice critical to the task (outbound approach to the L vs. R maze arm bifurcation, Fig. 1b, **Fig. S1b**), analysis was performed only for outbound direction-based cell samples (defined above; inbound cell samples yielded a similar result). Furthermore, two inclusion criteria were imposed to ensure accurate estimates: first, data was only considered from positions beyond the choice boundary (i.e. the L or R arm); second, only cell samples with at least 100 spikes in the L or R arm were considered.

The direction selectivity index was calculated for direction-based cell samples that had at least one place field in either the O or I direction.

The selectivity index was defined as (fr_2_ – fr_1_)/(fr_1_+fr_2_), where fr is firing rate and 1, 2 correspond to two alternative conditions: for location, 1: L, 2: R; for direction: 1: O, 2: I. For single-cell surveys in **Fig. S1h, k**, fr was defined as the peak firing rate from the time-averaged spatial firing maps (see above); in other analyses, mean firing rate is also used, and stated explicitly where the case.

### Theta cycles and theta phase

For each subject, a tetrode in CA3 yielding clustered putative principal cells (i.e. located in the principal cell layer) was designated as the LFP recording site. In a minority of recording epochs (30 out of 287) for which a tetrode in the principal cell layer of CA3 was not available, a tetrode located in principal cell layer of CA1 was used instead.

To isolate activity in the frequency range of hippocampal theta^27,30^, LFP from these recording sites was filtered at 5–11 Hz. Peaks and troughs of the filtered LFP were detected and used to define half-cycles (π radians) by linear interpolation^32,129^. To establish a common reference phase, a phase histogram (π/6 or 30° bin size) of aggregate single (principal) cell firing in CA1 was calculated across locomotor periods for each recording day; the phase of maximal CA1 firing was then assigned to be 0°^44^, with the half-cycle offset (±π) corresponding to the phase segregating individual cycles. Theta cycles were identified as individual cycles whose duration was consistent with the 5-11 Hz frequency range (<200 ms (5 Hz) and >90 ms (11 Hz)); intermittent cycles not meeting this criterion were disregarded in subsequent analyses that explicitly reference theta cycles or theta phase.

### Theta phase locking

To survey the prevalence and strength of theta rhythmicity in neural firing, theta phase histograms of single-cell firing were calculated. For a given cell, theta phase locking analysis was performed for locomotor periods (>4 cm/s), and moreover only when at least 50 spikes were available. Firing was combined across all available maze epochs. For CA2 (**Fig. S1m**, middle column), cells previously classified as N cells (see above) were excluded from this analysis, as theta-locking measures for these cells in the present dataset have been reported previously^22^.

### Firing correlograms

To identify temporal patterns in neural activity, firing correlograms (CG) (time histograms; ±1.5 s, 5-ms bins) were calculated for single cells (auto-correlograms, ACG; t = 0 bin set to 0) and cell pairs (cross-correlograms, XCG). Data analyzed was restricted to locomotor periods that lasted at least 1.5 s, with further subdivisions detailed below. When multiple epochs for a maze environment were available, data was pooled across epochs. If data from two unique maze environments were available, then data from the two unique maze environments were analyzed independently; thus a cell pair (XCG) or single cell (ACG) could contribute more than one CG.

Two types of CGs were calculated: count and corrected. Count CGs were calculated by summing CGs (spike counts) across all data periods. For corrected CGs, CGs from each individual data period were first triangle-corrected^30,130^ to offset bias due to data periods of variable lengths (corrected spike counts); the corrected CG was then obtained by taking the mean over all individual data period CGs.

CGs (count and corrected) were then processed at two timescales: coarse (±1 s window) and fine (±0.4 s window). At each timescale, CGs were convolved with Gaussian kernels of an appropriate bandwidth (coarse, σ = 50 ms; fine, σ = 10 ms; meant to capture behavior- and theta-timescale activity, respectively^107,131,132^) and then peak-normalized within the respective (coarse or fine) time window. Processed count CGs are shown in single example plots (e.g. Fig. 1e, h) and survey plots (XCG: Fig. 1e, **Fig. S3f-h**; ACG: Fig. 2f) (example raw count CGs shown in **Fig. S3e** for illustration), while processed corrected CGs were used for subsequent analyses. Both plots and analyses were restricted to CGs with at least 100 spikes in the fine timescale window (±0.4 s). Cycling firing at 8 Hz (“skip” firing^33,34,133^) was overtly present in both count and corrected CGs.

CGs were calculated from locomotor period data subdivided by either (I) path (II) direction, or (III) choice condition. As with other analyses in this study, locomotor periods were defined as periods of movement speed >4 cm/s; a threshold of >20 cm/s was also used to evaluate whether a firing pattern of interest (cycle skipping, see below) required low movement speed (lower panels in **Fig. S4**).

(I) For path, CGs were calculated from data separated by periods corresponding to the four task path types; thus a given cell (or cell pair) in a given maze environment could contribute up to four samples in subsequent path-based analyses (path-based cell (or cell pair) samples). Path-based CGs were used to survey firing in the dataset (ACG: Fig. 2a-f, XCG: **Fig. S3h**) and to compare differences between recording regions (Fig. 2g, **Fig. S4a, c**); subdividing data by path type distinguishes both path- and direction-dependence^26,47,50,108,109^ of hippocampal place cell firing, and in this way maximally distinguishes between putatively independent firing correlates. The finding of significant differences in firing between recording regions (Fig. 2g or **Fig. S4a**, **c**) was also observed without subdividing data or by subdividing by direction (data not presented).

(II) For direction, CGs were calculated from data separated by periods corresponding to outbound vs. inbound periods; thus a given cell (or cell pair) in a given maze environment could contribute up to two samples in subsequent path-based analyses (direction-based cell (or cell pair) samples). Direction-based XCGs are presented in the cell pair examples of Fig. 1, **Fig. S1n-q, Fig. S2**, **Fig. S3a-e**, and also surveyed across all cell pairs in **Fig. S3f, g**. Direction-based ACGs (data not presented) were qualitatively similar to path-based ACGs (Fig. 2e, f).

(III) For choice condition, CGs were calculated from data (a) recorded from linearized positions within the choice boundary, and (b) subdivided into outbound vs. inbound periods; given (b), a given cell in a given maze environment could contribute up to two samples in subsequent analyses (choice-based cell samples). Choice-based CGs were subsequently analyzed (cycle skipping analysis, see below; ACG: Fig. 2h, **Fig. S4b**, XCG: **Fig. S4d**) to assess whether choice behavior (choice passed: inbound samples; choice imminent: outbound samples) was a correlate of temporal firing patterns.

### Cycle skipping index (CSI)

Cycling firing at 8 Hz was detected and quantified with a theta cycling index (CSI) that was conceptually equivalent to a previously described theta “skipping” index^33,34^; the goal of either approach is to measure the lack of firing on adjacent theta (8 Hz) cycles. In the present study, the term “cycling” was adopted to refer explicitly to the observation that periods of “skipping” (i.e. lack of firing) in the firing of one group of cells can correspond to periods of firing in another group of cells, furthermore in the case where the two groups encode mutually exclusive scenarios (suggested initially at the cell-pair level, Fig. 1, **Fig. S2**).

Two types of CSI were calculated: one for single cells (cell CSI) and one for cell pairs (cell pair CSI); for both, the calculation was performed on corrected CGs (ACG for cell CSI; XCG for cell pair CSI) that had at least 100 spikes within the fine timescale window (±0.4 s) and that were smoothed and peak-normalized (see above).

#### Single-cell cycling (cell CSI)

For each ACG, two local maxima (peaks: p_1_ and p_2_) within two respective time windows were detected: p_1_, the peak nearest t = 0 in the 90-200 ms window; p_2_, the peak nearest t = 0 in the 200-400 ms window. In some cases, a peak was not detected within a time window: for p_1_, the maximum value in the window was then used; for p_2_, the minimum value in the window was then used.

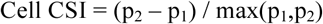

For two ACGs (path-based samples), the CSI could not calculated due to an absence of spiking in both the 1^st^ and 2^nd^ peak time windows.

#### Cell-pair cycling (cell pair CSI)

For each XCG, five local maxima (peaks: p_0_, p_±1_, p_±2_) within five respective time windows were detected. First, p_0_, the peak nearest t = 0 in the ±90 ms window was identified (if no peak was detected, then the p_0_ was set as the value at t = 0). If p_0_ did not occur at t = 0, then the four remaining peaks were detected in time windows relative to the time bin of p_0_. The four remaining peaks were defined as follows: p_±1_, the two peaks nearest t = 0 in ±90-200 ms windows; and p_±2_, the two peaks nearest t = 0 in ±200-400 ms windows.

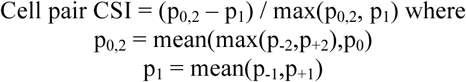

### Cell pair firing types

Cell pair samples with at least 100 spikes in ±0.4 s of XCG were classified into one of four types: (1) classic, (2) anti-synchronous cycling (anti), (3) super-synchronous cycling (super), and (4) off cycling (off). Classic corresponds to the pattern of co-firing expected given single-cell firing on adjacent cycles (**Fig. S1n-q**), as previously described in the hippocampus^44,107^. Anti corresponds to a pattern of co-firing in which the two cells consistently fire on alternate 8 Hz cycles; Super corresponds to a pattern of co-firing in which the two cells consistently fire together every other 8 Hz cycle. Off corresponds to a pattern of co-firing showing either relatively more or relatively less firing every other 8 Hz cycle, yet for which synchronization within 8 Hz cycles was ambiguous.

Formal criteria were as follows: classic pairs were those with cell pair CSI <0.3; anti cell pairs were those with p_0_ (see above) detected within the ±40 ms window in the XCG, and had cell pair CSI <-0.3; super cell pairs were those with p_0_ detected in ±40 ms in the XCG, and had cell pair CSI >0.3; off cell pairs were all other cell pairs that had abs(cell pair CSI) >0.3.

The primary goal of these criteria was to detect cell pairs that unequivocally exhibited cycling firing (**Fig. S3e**, **Fig. S3f-h**) rather than to demarcate cell types; from informal observations, the cell pair CSI cutoff value presently chosen (0.3) tends to over-classify cell pairs as classic. Given the adopted criteria, cell pair proportions were tabulated (**Fig. S3f-h**), moreover when cell pairs recorded from the same tetrode were excluded (yielding similar proportions; see caption of **Fig. S3f-h**).

It is important to note that the present analysis approach assumes that cycling dynamics do not exceed two cycle types (A-B-A-B-…), though it is possible that cycling dynamics occur with three or more cycle types. Given this limitation, single-cell measures (Fig. 2; **Fig. S4a-b**) were adopted to detect cycling dynamics; the cell pair results are presented here as an initial approach to the observation of constant cycling dynamics in the hippocampus (Fig. 1, **Fig. S2, Fig. S3**), and as a parallel to recent results in entorhinal cortex^33,34^.

### Clusterless decoding

To evaluate neural representation at the population level, Bayesian decoding of unsorted neural spikes (i.e. unassigned by experimenter to single cells) was performed^40,41^. The inclusion of unsorted spikes is advantageous for population-level analysis in that recorded data subject to analysis is maximized: all spike sources monitored by electrodes are analyzed. Furthermore, recent studies report improved decoding performance for hippocampal data^40,41^. This improvement is deducible given that (i) spikes with similar waveforms emanate from the same cells^111,134,135^, a correspondence not dependent on spike sorting, and (ii) multi-tetrode recording in hippocampus routinely yields a substantial number of high-amplitude spikes left unsorted prior to analysis^49^. In the present study, two variables were separately decoded: location^13,104,136,137^ and heading direction^26,49,50,106^, each previously established as single-cell representational correlates in the hippocampus^138^.

#### Data criteria

To limit analysis to population-level activity, decoding was performed only for recording epochs for which there were at least 20 putative principal cells clustered and firing at least 100 spikes, moreover only for subjects with at least three such epochs available (83 epochs across 7 subjects). Within each epoch, spikes included for analysis were required (i) to exceed 60 µV on at least one tetrode channel (ii) to be recorded on a tetrode that yielded at least one clustered putative principal cell. Across qualifying epochs, 4-17 (median: 9) tetrodes per epoch met criterion (ii); these tetrodes were predominantly in CA1 and CA3, with a subset of epochs (34 out of 83 epochs) also including tetrodes in CA2 and DG. Restricting decoding to CA1 and CA3 tetrodes yielded qualitatively equivalent findings (not presented). In example plots (e.g. Fig. 3a-c, Fig. 5a-c), spikes analyzed were aggregated across tetrodes and shown as multi-unit activity (MUA).

#### Analysis times

For both location and direction, the decoding procedure was performed within each ~15 min epoch; data used to construct an encoding model were from locomotor periods while decoded data was from locomotor periods and flanking 0.5 s periods (stopping periods). Flanking stopping periods were included in the analysis since the theta rhythm and associated neural firing are known to continue to be expressed during these times; these periods were also included to assess neural activity possibly associated with head scanning behaviors^11,12^, which can encompass brief low-speed periods. In plots, neural activity overlapping with stopping periods are indicated (overlaps stop; Figs. 3d-e, Fig. 5d-e, **Fig. S6g-h**).

For location, spikes analyzed were further restricted to outbound periods to eliminate the contribution of direction-specific activity to the encoding model. For direction, spikes analyzed were not restricted by direction, but rather restricted to periods when subjects were located in the three parallel longer segments of the maze, specified as linearized positions (defined with respect to the center reward well; see above) that were either less than or more than 40 cm from the linearized position of the central junction (shown as lighter grey positions in example behavioral maps in Fig. 5a-c, **Fig. S10b-e**). This restriction was enforced to facilitate comparison across studies; in particular, prior work on hippocampal directional coding in mazes^26,50,106,139^ has referred to straight rather than jointed (e.g. perpendicularly connected) maze segments.

#### Decoding time windows

For location, decoding windows were 20 ms with 4 ms overlap between windows. For direction, two types of decoding windows were used (illustrated in **Fig. S9b-c**): (i) 20 ms with 4 ms overlap (sliding window decoder), and (ii) windows correspondent with theta half-cycles (half-cycle decoder; Fig. 5, **Fig. S10a-d**). For (ii), theta phase estimated from LFP (see above) was used to identify windows of duration π/4 (90°) centered on first and second halves of theta (i.e. first-half window: (−3*π/4, -π/4); second-half window: (π/4, 3*π/4)). These windows were chosen on the basis of results at both the single-cell (Fig. 4e-h) and population (**Fig. S9d**) levels indicating that representation of non-current (hypothetical) direction is weakest and strongest at approximately –π/2 and π/2, respectively.

It is worth noting that decoding is performed in sub-second time windows (20-50 ms) to assess temporal dynamics; in contrast, the encoding model is constructed from data pooled across the ~15 min recording epoch, moreover without referencing temporal dynamics in the epoch.

#### Algorithm

A two-stage decoding algorithm described previously^40,41^ was used. In the first stage (encoding), the mapping between spikes and the representational variable (location or direction) was modeled as an N-dimensional probability distribution (mark space, M), where each spike corresponds to an N-dimensional vector (mark). In M, N-1 dimensions correspond to the feature space of spikes while the remaining dimension corresponds to the representational variable. M is estimated from all spikes occurring during encoding periods using kernel density estimation. In the present case, N is 5, where 4 dimensions correspond to the amplitude (µV) of the spike on each of the spike’s 4 parent tetrode channels while the remaining dimension corresponds to the value of the representational variable (S; location: linearized position (cm); direction: −1 for inbound, 1 for outbound) observed from the subject at the time of the spike. Each spike contributed a 5-D Gaussian kernel, with the four amplitude dimensions contributing symmetrically (each σ = 20 µV), symmetric in the amplitude dimensions). For location, the representation dimension of M was linearized position divided into 1-cm bins, with each spike contributing a Gaussian kernel (σ = 8 cm). For direction, the representation dimension of M was two-bin distribution, with each spike contributing a Kronecker delta kernel.

In the second stage, decoding was performed using Bayes’ rule:

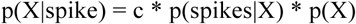

where spike refers to the set of spike marks observed in the decoding window, and c is a normalization constant. Each p(•) term is a probability density over the representational dimension of M; p(X|spikes) is the posterior (decoded output; estimate of the representational correlate X); p(spikes|X) is the likelihood (encoding model); p(X) is the prior.

To obtain p(spikes|X), the aggregate spiking activity across tetrodes was modeled as a marked point process with mark space M, with spikes in each decoding time window treated as independent and following a Poisson distribution with rate parameter fully characterized by M (derivation and formalism in earlier report^41^).

Next, the prior p(X) was taken to be either uniform or history-dependent. For decoding of direction, a uniform p(X) was used given a lack of prior knowledge of population-level representation of direction in the hippocampus. For decoding of location, both uniform^13,21,40^ and history-dependent^11,136,137,140^ priors were used to assess the generality of the population-level result (constant cycling at 8 Hz between alternative locations; history-dependent prior: Fig. 3; both priors: **Fig. S6**). The advantage of a uniform prior is that it minimizes assumptions about the decoded activity; the advantage of a history-dependent prior is that it models known properties of the decoded activity.

Taking previous work^11,140^ as a starting point, the history-dependent prior was designed to model the observation that population-level spiking in the hippocampus regularly encodes locational sequences that evolve at virtual speeds exceeding 1 m/s^42,49,141,142^. This property can be captured by a Markovian state-space model^41^ implemented by a prior defined at each time step as the product of a constant 1-step transition matrix and the posterior from the previous time step. To eliminate bias for the decoded output to evolve in a particular direction, state transitions were modeled as a 2D random walk^136,137,143^ where, for simplicity, the three arms of the maze were treated as locations in a radially symmetric Y shape (120° between arms). Transition probabilities between maze locations were calculated as the value of a Gaussian (σ = 1 cm) evaluated at the Euclidean distance between each location in the Y. To model virtual speeds exceeding 1 m/s, the transition matrix was exponentiated by 10^11,140^; thus for decoding windows that shifted every 4 ms, the 1-cm scale of the Gaussian corresponds to a virtual speed of 2.5 m/s, with exponentiation by 10 corresponding to 25 m/s. The resulting prior is approximately an order of magnitude more conservative (diffuse over spatial locations) compared to priors modeling virtual speeds of ~2-5 m/s^11^. It is also worth noting that since the model (transition matrix) is constant, it cannot impose rhythmic patterns (e.g. continuous 8 Hz rhythmicity).

### Constant cycling of location

To determine whether constant cycling at 8 Hz between alternative locations (i.e. left (L) vs. right (R) maze arm; observed initially in cell pairs (Fig. 1)) occurred at the population level, the output of the clusterless decoding of location (decoded posteriors; see above) was analyzed. Moreover, to investigate further the finding of elevated cycling dynamics when subjects were approaching the L vs. R spatial choice (Fig. 4h, suggesting cycling between L vs. R locations), analysis was restricted to the three types of maze passes (defined above) entailing this behavior: outbound Left path passes, outbound Right path passes, and backtrack passes from the center well. Analysis was further restricted to locomotor periods and flanking stopping periods (0.5 s), to periods when the subject was located within the choice boundary (center zone), and to periods when the subject was located at least one-third of the linear distance to the choice boundary along the center arm; the latter two restrictions were imposed to focus analysis on cycling neural activity associated with choice approach (initially identified at the single-cell level (Fig. 2h, **Fig. S4b, d**).

First, 8 Hz cycles were segregated on the basis of the ~8 Hz theta rhythm. Prior results suggest that population-level representation of locational sequences occur within individual theta cycles^42-44,49,142^: thus the initial step was to identify an appropriate phase by which to segregate theta cycles. To this end, the theta phase distribution of the representation of alternative locations (L or R maze arm) was calculated for each recording epoch; specifically, the decoded probability density located in either the L or R arm (integrating over position bins) was histogrammed in 30° theta phase bins (linear interpolation of 5-11 Hz LFP, see above). The phase bin having the minimum probability density was then used to segregate theta cycles.

Second, candidate 8 Hz cycles representing the alternative locations (L or R maze arm) were identified and binarized. Candidate cycles were identified as the segregated theta cycles that had probability density >0.1 for either the L or R maze arm (L/R density; the remainder corresponding to the center arm); then, for each candidate cycle, the arm with the higher probability density was designated as the location (L vs. R) represented.

Third, constant cycling periods were detected. In general, constant cycling can be defined as cycle-to-cycle switching of representation across at least 3 successive cycles, i.e. A-B-C-D where B is not A, C is not B, and D is not C. For binary path choice (between L vs. R; **Fig. S1b**), constant cycling is defined only for the special case of A-B-A-B; accordingly, putative constant cycling periods were detected as cases where the representation switched for at least 3 successive cycles (L-R-L-R… or R-L-R-L…), corresponding to a minimum total duration of 4 cycles. The start and end of single constant cycling periods were defined as the beginning of the first cycle to the end of the last cycle. Note that constant cycling periods were only detected within periods of contiguous candidate theta cycles (contig period).

To evaluate whether constant cycling could have resulted from random activity or noisy data, the observation of constant cycling was tested against a null model in which cycle order was random (temporal shuffle); testing was conducted at the study-wide level and for individual constant cycling periods.

At the study-wide level (Fig. 3d, **Fig. S6g**), a P-value was calculated by randomly shuffling (10000 permutations) the order of all candidate theta cycles within every contig period across all recording epochs. For each shuffle, constant cycling periods were then re-detected, after which the total number of cycles participating in the re-detected constant cycling periods was tabulated; the P-value was the proportion of shuffles for which the total number of such cycles (i.e. those in constant cycling periods) was equal or greater to the number in the observed data.

For individual constant cycling periods (Fig. 3e, **Fig. S6h**), a P-value was calculated for each observed constant cycling period by randomly shuffling (10000 permutations) the order of candidate theta cycles within the same recording epoch and within time periods of the same path pass type (outbound Left, outbound Right, or center well backtrack) as that of the observed constant cycling period; the P-value of the constant cycling period was proportion of shuffles for which a constant cycling period of the same or greater cycle duration was detected within the same contig period as the observed constant cycling period. The P-value thus measures the frequency of the representational activity pattern (an individual constant cycling period) with respect to the empirical prevalence of the components of that pattern (L vs. R candidate theta cycles from the same recording epoch and path pass type). A criterion of P < 0.05 was then adopted for subsequent analysis of individual constant cycling periods (Fig. 3f, g, **Fig. S6i, j**).

### Constant cycling of direction

To determine whether constant (half-theta) cycling between directions (suggested by initial observations; Fig. 4, **Fig. 10a**) could occur at the population level, the output of the clusterless decoding of direction (decoded posteriors; see above) was analyzed. Analysis was conducted for all locomotor periods and flanking stopping periods (0.5 s). Furthermore, analysis was performed on the output of the theta half-cycle decoder, as single-cell and population-level results (Fig. 4e-h, **Fig. S9**) suggested that this decoder would be maximally sensitive to representations of alternative directions.

First, the decoded half-cycles were binarized (illustration in **Fig. S9b-c**); for each half-cycle, the direction with the higher probability density was designated as the direction represented (outbound (O) vs. inbound (I); see above).

Second, constant cycling periods were detected with a procedure analogous to that of location. In general, constant cycling can be defined as cycle-to-cycle switching of representation that occurs contiguously for least 3 successive cycles, i.e. A-B-C-D where B is not A, C is not B, and D is not C. For binary heading direction (O vs. I; **Fig. S1i**), constant cycling is defined only for the special case of A-B-A-B; accordingly, putative constant cycling periods were detected as cases where the representation switched for at least 3 successive theta half-cycles (O-I-O-I… or I-O-I-O…), corresponding to a minimum total duration of 4 theta half-cycles. The start and end of single constant cycling periods were defined as the beginning of the first theta half-cycle to the end of the last theta half-cycle. Note that constant cycling periods were only detected within periods of contiguous theta half-cycles (contig period).

Third, constant cycling periods were identified as occurring within three types of directional periods: inbound, outbound, or mixed directional periods. The first two types were identified if the detected constant cycling period occurred entirely within the respective (inbound or outbound) directional period. The mixed period type was identified if the constant cycling period overlapped with both inbound and outbound periods.

As in the case of location (see above), to evaluate whether constant cycling could have resulted from random activity or noisy data, the observation of constant cycling was tested against a null model in which cycle order was random (temporal shuffle); testing was conducted at the study-wide level and for individual constant cycling periods.

At the study-wide level (Fig. 5d), a P-value was calculated by randomly shuffling (10000 permutations) the order of all half-theta cycles within every contig period across all recording epochs. For each shuffle, constant cycling periods were then re-detected, after which the total number of half-theta cycles participating in the re-detected constant cycling periods was tabulated; the P-value was the proportion of shuffles for which the total number of such half-theta cycles (i.e. those in constant cycling periods) was equal or greater to the number in the observed data.

For individual constant cycling periods (Fig. 5e), a P-value was calculated for each observed constant cycling period by randomly shuffling (10000 permutations) the order of half-theta cycles within the same recording epoch and within the same directional period type (inbound or outbound) as that of the observed constant cycling period; the P-value of the constant cycling period was proportion of shuffles for which a constant cycling period of the same or greater cycle duration was detected within the same contig period as the observed constant cycling period; for individual constant cycling periods that occurred during mixed directional periods, shuffling was performed simultaneously and separately for the half-theta cycles that occurred respectively within outbound vs. inbound periods. The P-value measures the frequency of the representational activity pattern (an individual constant cycling period) with respect to the empirical prevalence of the components of the pattern (O vs. I half-theta cycles from the same recording epoch and directional period type). A criterion of P < 0.05 was then adopted for subsequent analysis of individual constant cycling periods (Fig. 5f, g). In addition, analysis of individual constant cycling periods that lasted at least 8 (half-theta) cycles was also conducted (**Fig. S10f, g**).

### Decoding choice

To assess whether population-level activity in the hippocampus could predict spatial choice (**Fig. S7**), Bayesian decoding of hippocampal neural firing was performed using the clusterless decoding algorithm (see above).

The decoding procedure (data criteria, analysis times, algorithm; see above) was the same as that of location and direction, but with the following specifications: (1) the data analyzed were from outbound path pass periods (left (L) and right (R); each pass treated as a single trial), (2) the representational variable was the path chosen (L vs. R in a two-bin distribution; −1: L path, 1: R path), (3) each spike during outbound path periods (occurring during either a L or R outbound path pass; see above for path period definition) contributed a Kronecker delta kernel to the two-bin choice dimension of the mark space (M; the encoding model), and (4) spikes used to encode and decode were restricted to the windows of duration π/4 (90°) centered on first and second halves of theta (first-half window: (−3*π/4, -π/4); second-half window: (π/4, 3*π/4)), respectively (half-cycle encoder and half-cycle decoder, respectively). Analysis was restricted to path passes (trials) that had at least 12 second-half windows available for decoding.

To evaluate different frameworks for interpreting hippocampal representation, three approaches to the decoding procedure were taken (described and schematized in **Fig. S7a-c**); each approach stipulates a specific subset of data for encoding and decoding.

The decoded output (posterior probabilities) was analyzed as follows. Within each trial, the probability density corresponding to the R choice was averaged across second-half (decoding) windows. If this average probability exceeded 0.5, then the decoded choice was taken to be R; otherwise, the decoded choice was taken to be L.

### Hypothetical representation: single cells

A body of work^29,138^ establishing that single neurons in the hippocampus encode externally observable variables was the basis of the present investigation into the neural representation of hypotheticals. Specifically, past findings indicate that single hippocampal neurons reliably (over single trials, recording epochs, days, and months^29,144,145^) fire more in specific conditions (e.g. a particular location, direction, trajectory, etc.) vs. other conditions (other locations, directions, trajectories), establishing that these cells encode the relevant variable (representational correlate). Methodologically, time-averaged tuning curves (e.g. place cell maps^13,29,104^) have been used to estimate single-cell encodings. This analysis approach to single cells was adopted, though with two differences.

The first difference was simply terminological: to make the underlying approach explicit (rather than the particular representational correlate, e.g. location, direction, etc.), the higher vs. lower firing conditions were generically termed the “preferred” (P) vs. “non-preferred” (NP) conditions, respectively.

The second difference was conceptual: in cases where single-cell firing was higher in the P (vs. NP) condition, firing in the NP condition was subsequently interpreted not as noise, but as possibly reflecting covert representation of the P condition, i.e. the non-current, or hypothetical, condition. Thus the goal of analysis was to quantify whether cells were tuned to fire more in one condition vs. others (P vs. NP) with the provisional interpretation that firing in the P vs. NP condition encodes current vs. hypothetical experience, respectively.

Data from each cell were analyzed as samples (cell samples) with the following criteria. If data from two unique maze environments were available for a cell, and the cell fired at least 100 spikes in the both environments, then the cell was analyzed in the two unique maze environments independently. Thus cells in the dataset could contribute more than one cell sample to subsequent analyses. If multiple recording epochs in one maze environment were available, then the recording epoch with the most recorded spikes was analyzed.

Four types of representational firing patterns were studied: (1) future path^46,47,109^ (previously termed prospective trajectory), (2) past path^46,47,109^ (previously termed retrospective trajectory), (3) directional^26,50^, and (4) a putative type termed “extra-field^11,146^”. As described above, path- and direction-based 1D firing maps for each cell sample were calculated.

For (1), direction-based firing maps were analyzed, moreover only the outbound direction; P vs. NP refers to L vs. R (or *vice versa*) outbound maze paths. For (2), inbound path-based maps (inbound L and inbound R) was analyzed; P vs. NP refers to L vs. R (or *vice versa*) inbound maze paths. For (3), direction-based (inbound (I) vs. outbound (O)) firing maps from each of the three maze arms were analyzed independently, with firing maps from each arm contributing independent cell samples if satisfying additional criteria specified below; for each cell sample, P vs. NP refers to the I vs. O heading direction, respectively (or vice versa). For (4), all four path-based maps were analyzed (outbound L, outbound R, inbound L, inbound R); P vs. NP refers to the task paths with vs. without at least one detected spatial firing field.

For each firing map, spatial firing fields (place fields), defined as contiguous linear positions of firing rate >2 Hz and at least 10 cm large, were detected separately in the following alternative conditions. For (1), (2), and (3), there were two alternative conditions: namely, for (1) and (2), L vs. R path, while for (3), O vs. I direction. For (4), there were four alternative conditions: outbound L vs. outbound R vs. inbound L vs. inbound R path. The inclusion criteria and identification of P vs. NP conditions for each type of representational correlate are as follows.

1. **Future path**. Place field detection was performed separately in two locational zones in the maze: the center zone (see above) and the outer maze arms (L and R). Only cell samples that had at least one place field detected in either of the outer arms (L or R), in addition to at least one place field detected in the center zone, were analyzed (note that since the location zones were contiguous, a place field detected in both zones could correspond to a single place field across the zones). Next, for the outer maze arms, the peak firing rate of all detected place fields were compared; the path type (L or R) corresponding to the maze arm (L or R) with highest peak firing rate place field was designated as the cell’s preferred (P) condition (e.g. L path), with the other condition designated as the cell’s non-preferred (NP) condition (e.g. R path).
2. **Past path**. Place field detection was restricted to the center arm. Only cell samples that had at least one place field for either path (L or R) were analyzed. The path type (L or R) with the highest peak firing rate place field was designated as the cell’s preferred (P) condition (e.g. L path), with the other condition designated as the cell’s non-preferred (NP) condition (e.g. R path).
3. **Directional**. Place field detection was performed independently in each maze arm. Only cell samples that had at least one place field for either direction (I or O) were analyzed. The direction (I or O) with the place field with the highest peak firing rate was designated as the cell’s preferred (P) condition (e.g. I direction), with the other condition designated as the non-preferred (NP) condition (e.g. O direction).
4. **Extra-field**. Place field detection was performed separately for each of the four task-based firing maps. Task paths with at least one detected place field were designated as the cell’s preferred (P) condition(s), while task paths for which no place fields were detected were designated as the cell’s non-preferred (NP) condition(s). Note that this definition of extra-field firing (i.e. firing in the NP condition) refers to path- and direction-specific firing, while previous approaches focus on location-specific firing^11,146^; the approaches are otherwise conceptually similar.

#### Selective firing (P vs. NP)

Past work indicates that the degree of path- and direction-specific firing exists on a continuum, with some cells firing almost exclusively in the preferred condition, while other cells fire equivalently in either condition (e.g. “bidirectional” cells or “pure” place cells). Since only cells showing differential firing between conditions can encode the relevant variable, it was therefore necessary to measure the selectivity of firing between conditions. To this end, the selectivity index (SI, defined above) was calculated for P vs. NP conditions; the SI was moreover calculated for both peak firing rate (from spatial firing maps) and mean firing rate.

A cell sample was classified as showing differential P vs. NP firing if two criteria were met: (1) the SI was >0.2 (equivalent to 1.5× higher firing rate in the P condition; other threshold values yielded qualitatively equivalent single-cell results) for both peak firing rates and mean firing rates, (2) the cell sample in the NP condition had no more than one detected place field. These criteria were adopted to limit analyses to cases where there was unequivocally higher firing on average in the P (vs. NP) condition.

### Theta phase of hypotheticals: single cells

To determine whether single-cell firing putatively encoding hypothetical experience (selective firing in preferred (P) vs. non-preferred (NP) conditions; see above) showed temporal organization at the sub-second timescale, neural firing in cells that were classified as showing differential firing was analyzed with respect to the phase of the ~8 Hz theta rhythm. Analysis was restricted to locomotor periods. In brief, spikes were subdivided by condition (P vs. NP) then analyzed separately.

The set of spikes analyzed was approximately the same as those used to construct firing maps, but with three differences: (1) a small proportion of spikes (~5%) were ignored if they did not occur within periods in which there was a valid estimate of theta phase (see above), (2) in the case of path firing (L vs. R), spikes were taken from all times during path passes between reward wells, with no exclusion of periods in which the rat was temporarily not within or oriented along the completed path, and (3) in the case of path firing (L vs. R), spikes were limited to those that occurred within the center zone. Criterion (2) was adopted to take an inclusive approach to the spikes analyzed, particularly given prior findings showing that path-specific firing generalizes to instances where the path is behaviorally interrupted^48^. Criterion (3) was adopted to restrict analysis to locations conventionally analyzed for path-specific firing (i.e. locations that overlap between paths).

Four representational correlates (see above) were analyzed: (1) future path, (2) past path, (3) direction, (4) extra-field. For each cell sample, the theta phases of the spikes in the P vs. NP conditions were then each used to construct separate theta phase histograms. Phase histograms were then constructed at two resolutions: 12-bin (30° bins) and 2-bin (180° bins; first and second half of cycle); the former (12-bin) solely for plots, while the latter (2-bin) for both plots and statistical testing of the basic hypothesis that representation in the hippocampus differs between the first and second halves of the theta cycle. Cell samples analyzed were restricted to those with at least 20 spikes in the phase histogram in the NP condition and with non-uniform phase histograms (Rayleigh tests at P < 0.05), in both the P and NP conditions.

For these cell samples, spike theta phases were averaged to obtain the mean angle in the P and NP conditions (Fig. 4b, f; first column in **Fig. S9a**). Second, phase histograms were normalized and averaged across cell samples (Fig. 4c, d, g, h; second and fourth columns in **Fig. S9a**). To evaluate whether firing was enhanced on the 2^nd^ half of the theta cycle, two comparisons were performed for the 2-bin phase histograms: (a) the proportion of firing in the 2^nd^ half of NP condition vs. 0.5 and (b) the proportion of firing in the 2^nd^ half of the NP condition vs. the 2^nd^ half of the P condition.

### Theta phase of hypotheticals: population-level

To determine whether representation of current vs. hypothetical experience showed temporal organization at the population level, the output of the clusterless decoding algorithm (see above), for both location and direction, was analyzed with respect to the phase of the ~8 Hz theta rhythm. Analysis was performed for locomotor periods. The analysis was analogous to the single-cell analysis (see above) in that theta phase was quantified separately (i.e. separate theta phase histograms) for different conditions; the preferred (P) vs. non-preferred (NP) conditions at the single-cell level correspond, here at the population-level, to current vs. hypothetical conditions, defined as follows.

The current condition is defined as the actual (veridical) state of the subject for a given representational correlate (e.g. the subject’s actual location if the representation is self-location), while the hypothetical condition is defined as the non-current state of the subject (e.g. a location remote from the subject’s actual location, or the direction opposite to that of the subject’s actual direction). The analysis procedure for each representational correlate studied presently (location and direction) is described in turn.

#### Location

To evaluate theta phase organization of the locational representation (Fig. 3h, **Fig. S6k**), the output of the clusterless decoding of location was analyzed for all outbound path passes (n = 1683 path passes across 7 subjects) when the subject was located in center zone. Two types of current condition were defined: current maze arm (Center) and the maze arm (L or R) chosen by the subject in the path pass (Choice, Fig. 3h); the alternative condition (Alt) was defined as the maze arm (L or R) not chosen by the subject in the path pass. Theta histograms were pooled across all decoding windows occurring within each path pass. Path passes were moreover analyzed inclusively, with no exclusion of times in which the subject was temporarily not within or oriented along the completed path; this inclusive approach was adopted given prior findings showing that path firing can generalize to passes where the path is behaviorally interrupted^48^.

#### Direction

To evaluate theta phase organization of the directional representation (**Fig. S9d**), the output of the clusterless decoding of direction (sliding window decoder; see above) was analyzed for each of the 7 subjects for which decoding was performed. The current condition was the subject’s current heading direction (outbound (O) or inbound (I)), while the hypothetical condition was the other direction (O or I). Theta histograms were constructed from decoding windows pooled across all epochs for each subject.

#### Phase histograms

Theta phase histograms (current and alternative/hypothetical) were calculated by first identifying the theta phase at the center time (average of start and end time) of each decoding window. Then, for each decoding window, the posterior probability density corresponding to each condition (current and alternative/hypothetical) was added to the correspondent theta phase histogram.

Phase histograms were constructed at two resolutions: 12-bin (30° bins) and 2-bin (180° bins; first and second half of cycle); the former (12-bin) solely for plots, while the latter (2-bin) for both plots and statistical testing of the basic hypothesis that representation differs between the first and second halves of the theta cycle. Phase histograms were then normalized within condition (current vs. alternative/hypothetical) and averaged across samples (passes or subjects). To evaluate whether representation was recruited on the 2^nd^ half a theta cycle, two comparisons were performed for the 2-bin phase histograms: (a) the proportion of firing in the 2^nd^ half of NP condition vs. 0.5 and (b) the proportion of firing in the 2^nd^ half of the NP condition vs. the 2^nd^ half of the P condition.

### Statistics

All statistical tests were two-sided.

### Code availability

All custom-written code is available upon request.

