## Supplemental figures for "Constant sub-second cycling between representations of possible futures in the hippocampus"

### Fig. S1. Study basics: localizing vs. generative, maze task, and established findings.

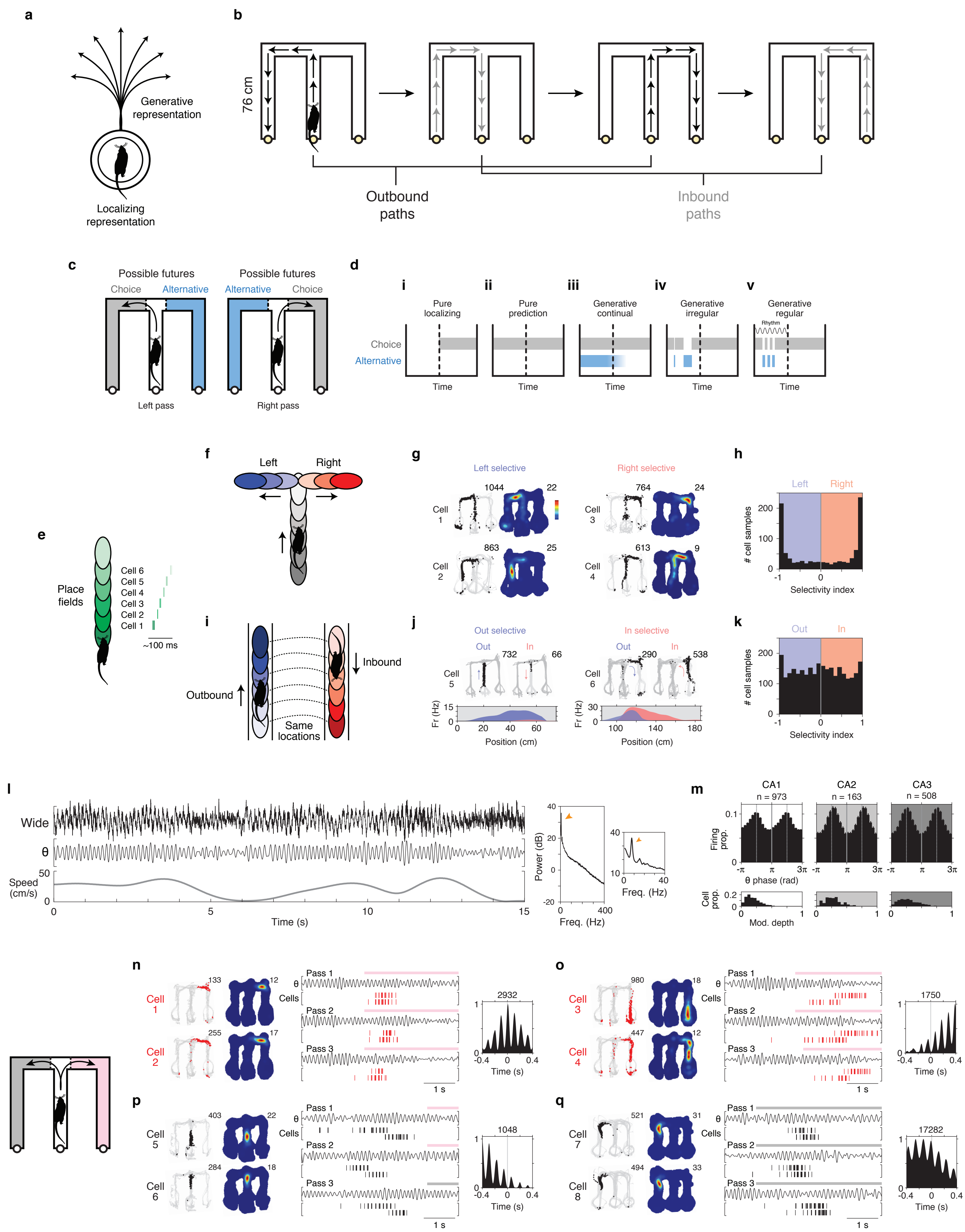

### Fig. S2. Cycling firing at 8 Hz: examples.

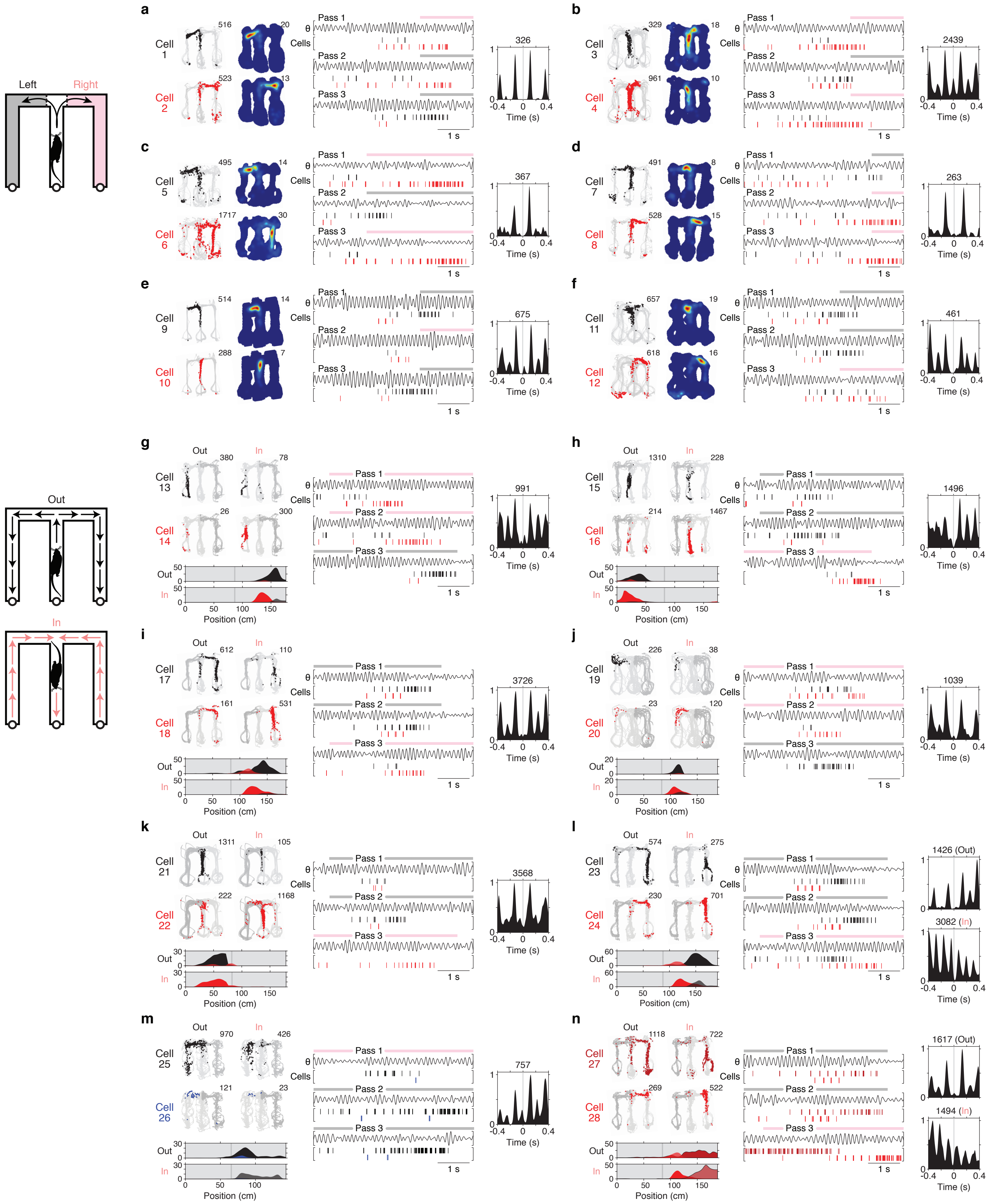

### Fig. S3. Cycling firing at 8 Hz: examples and survey of cell pairs.

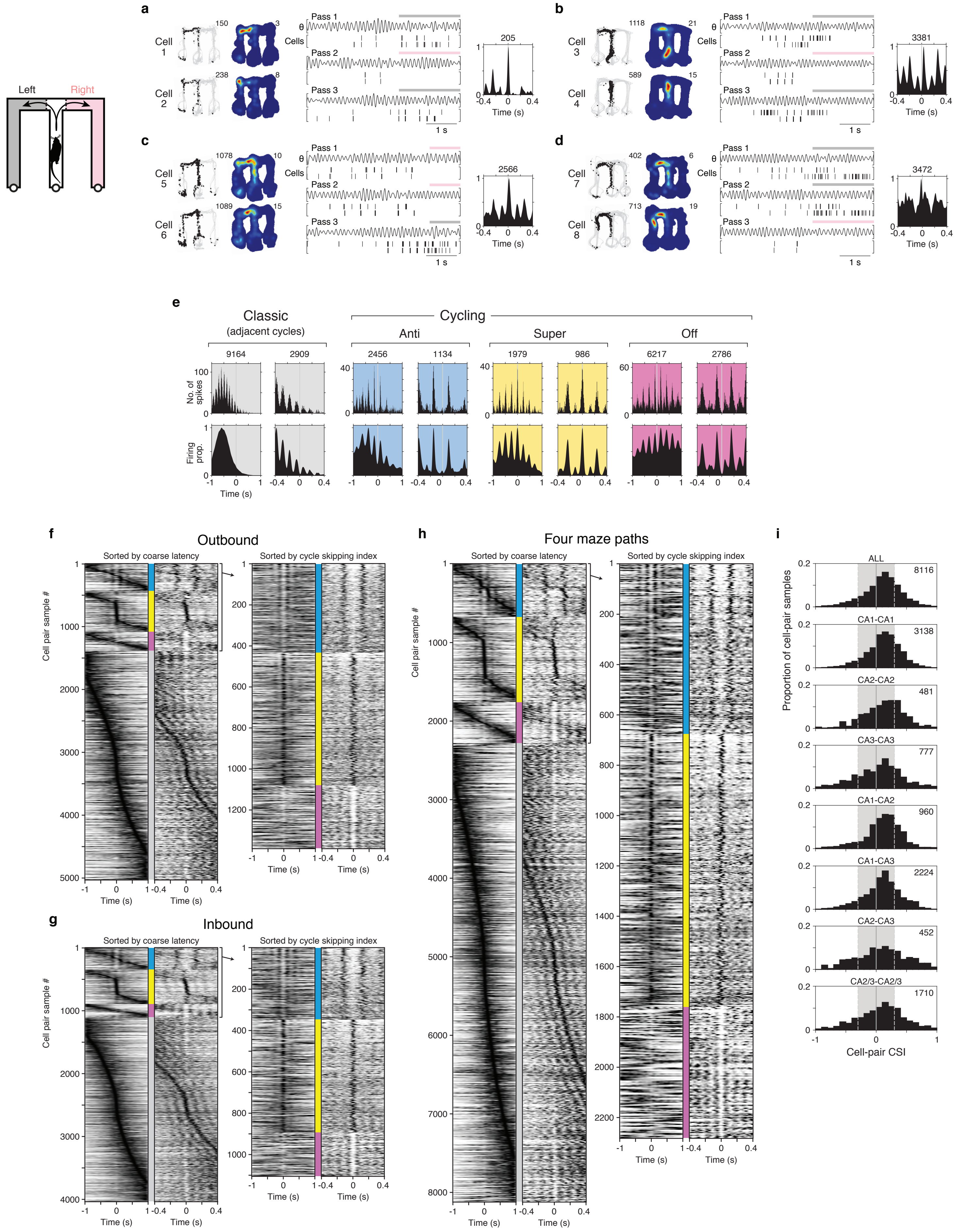

### Fig. S4. Cycling firing at 8 Hz: two correlates.

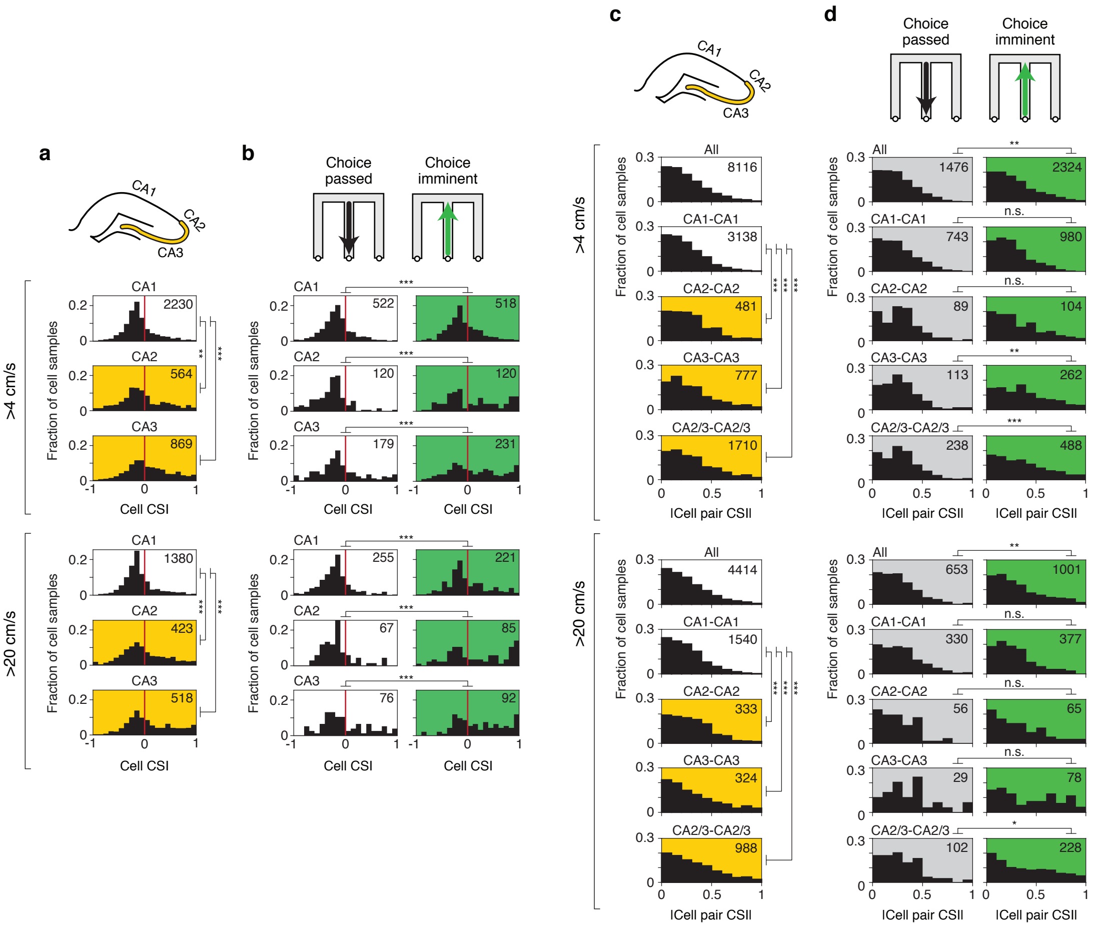

### Fig. S5. Constant cycling (8 Hz) at the population level: basic observation.

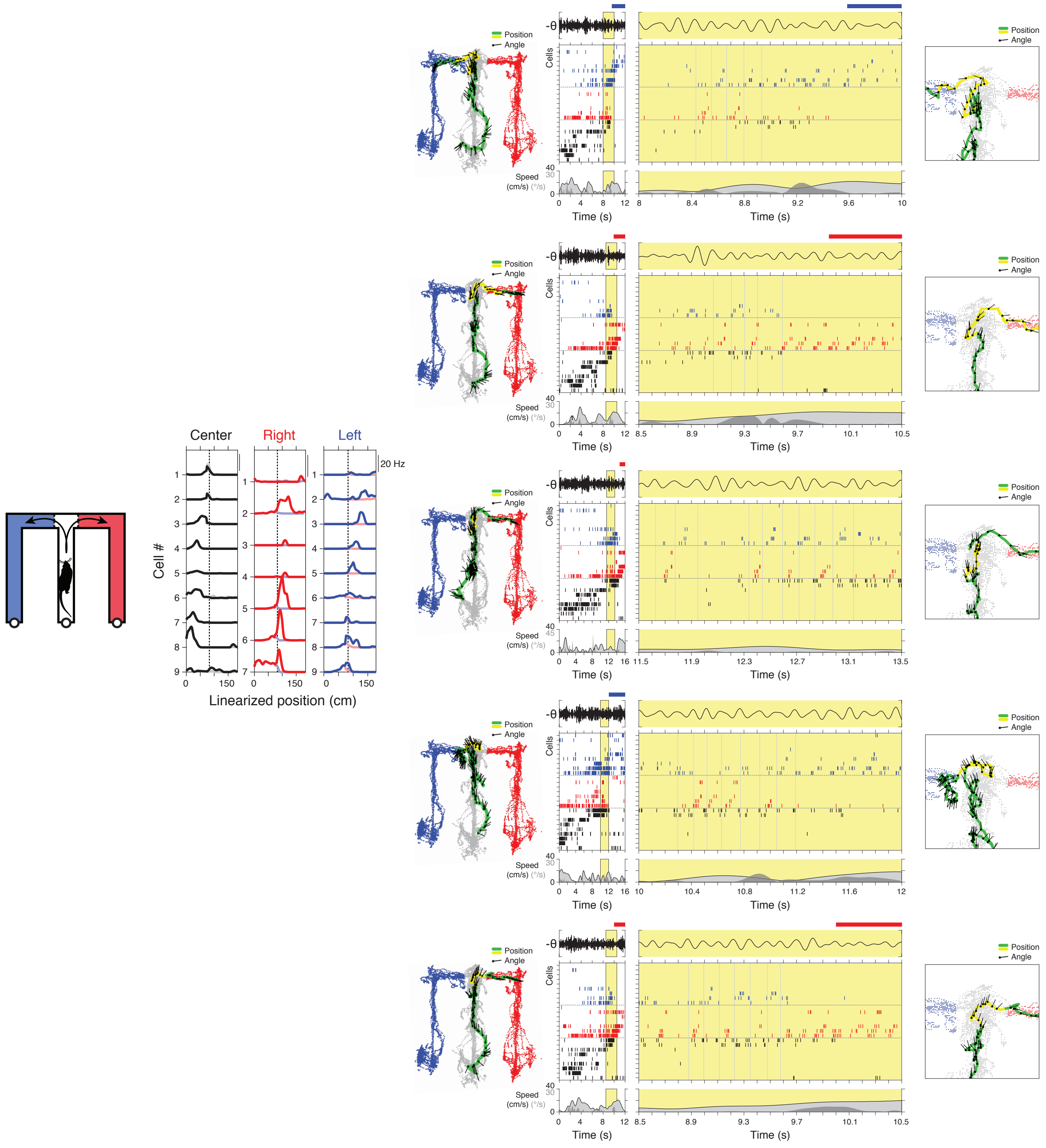

### Fig. S6. Constant cycling (8 Hz) between possible future locations: additional examples and approach.

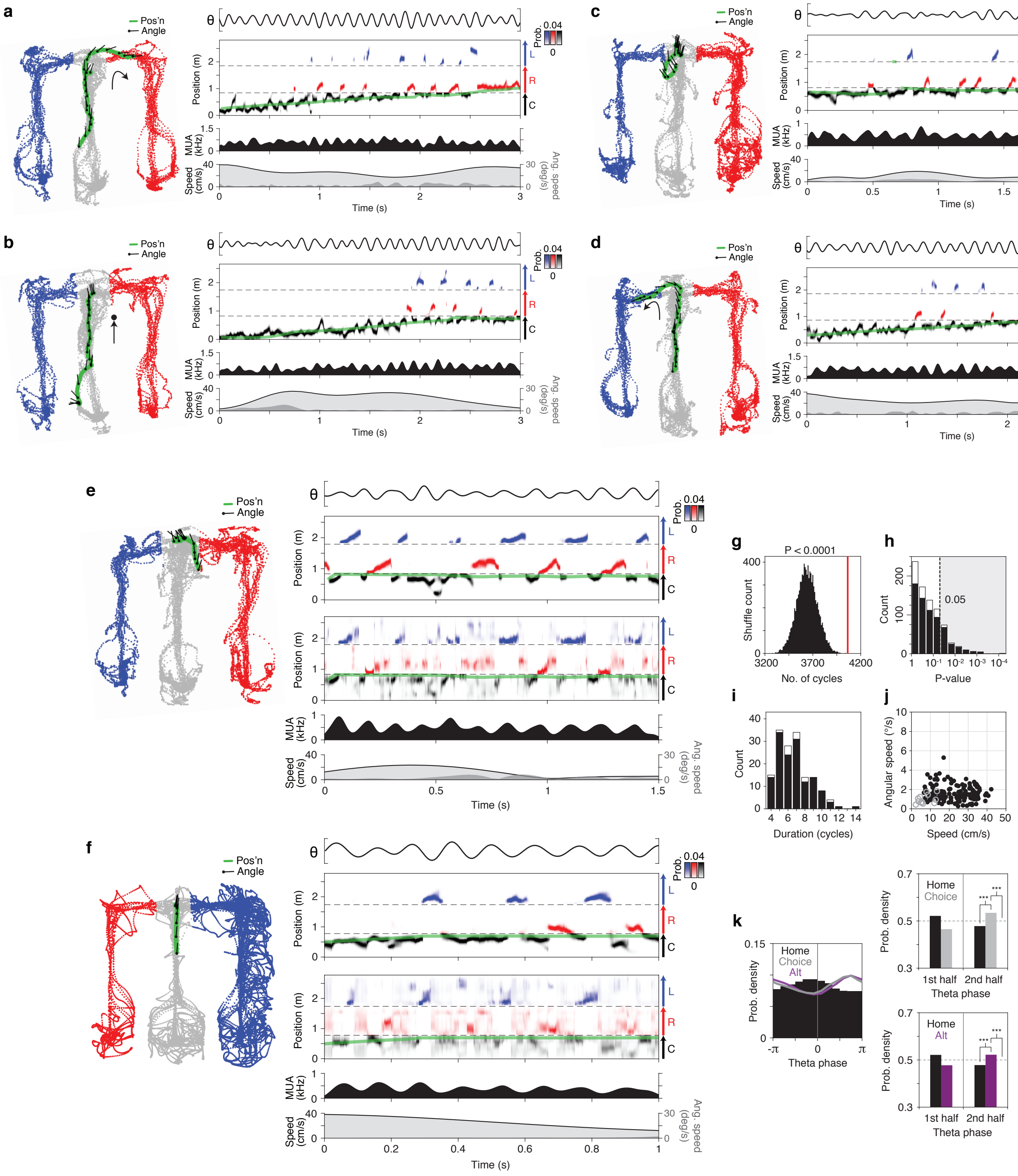

### Fig. S7. Decoding choice.

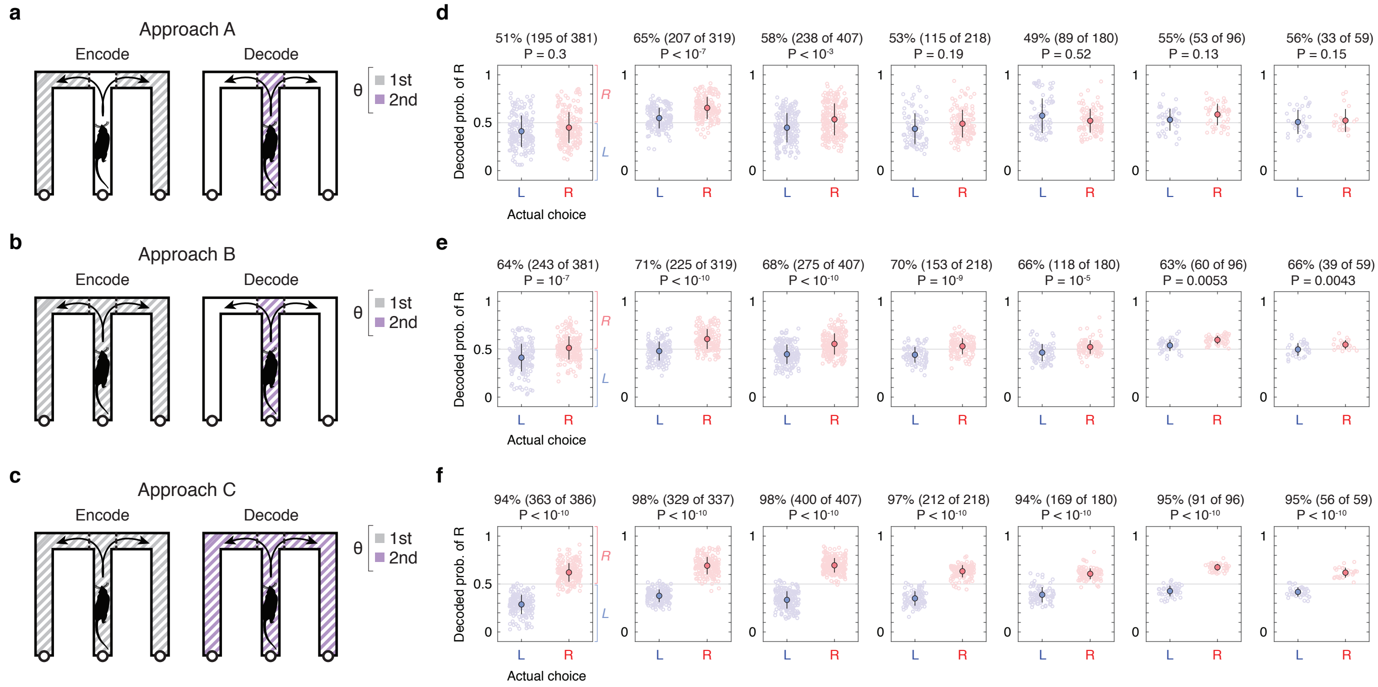

### Fig. S8. Intra-cycle coding of hypotheticals: additional examples.

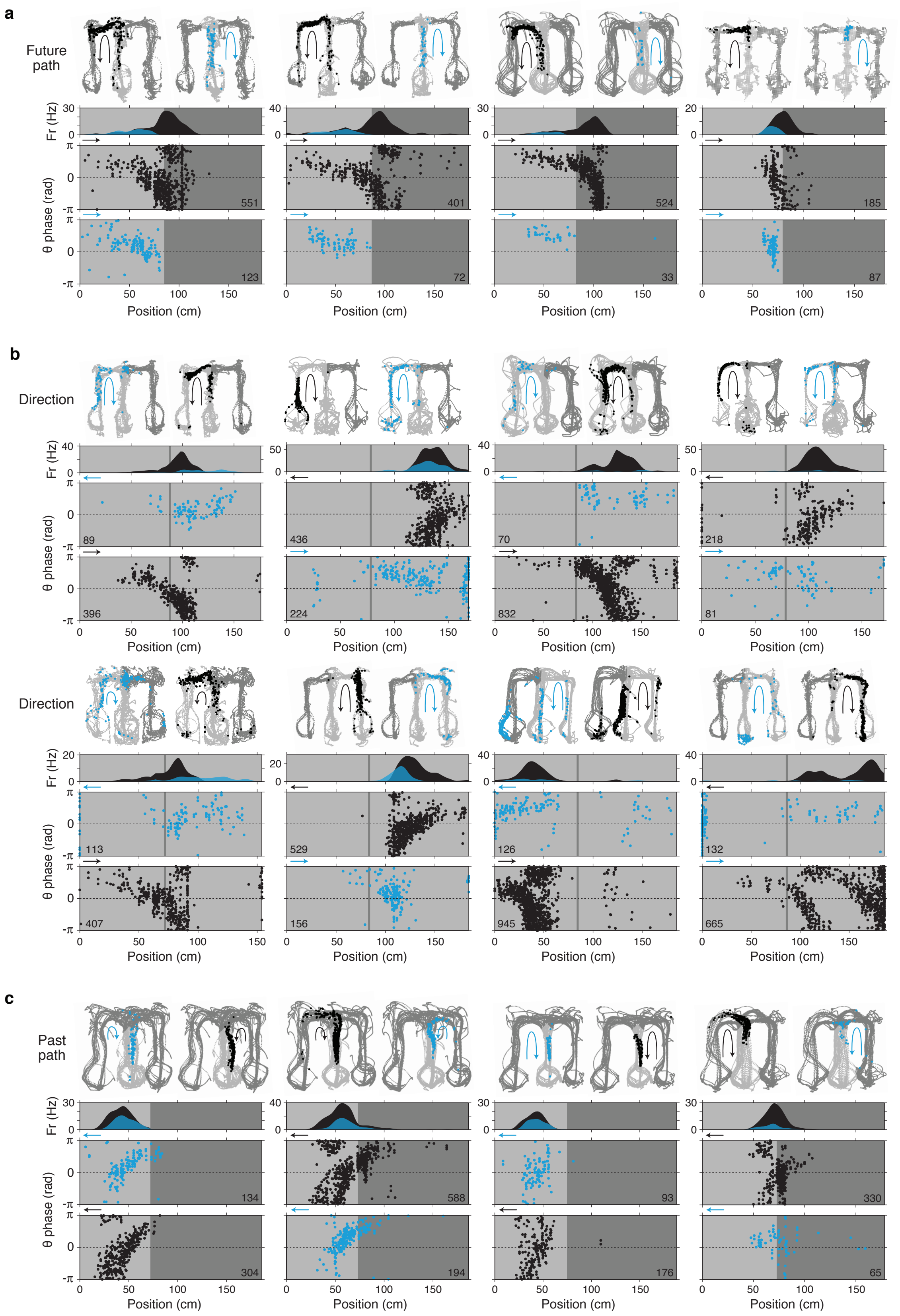

### Fig. S9. Intra-cycle coding of hypotheticals: single-cell summary and population decoding.

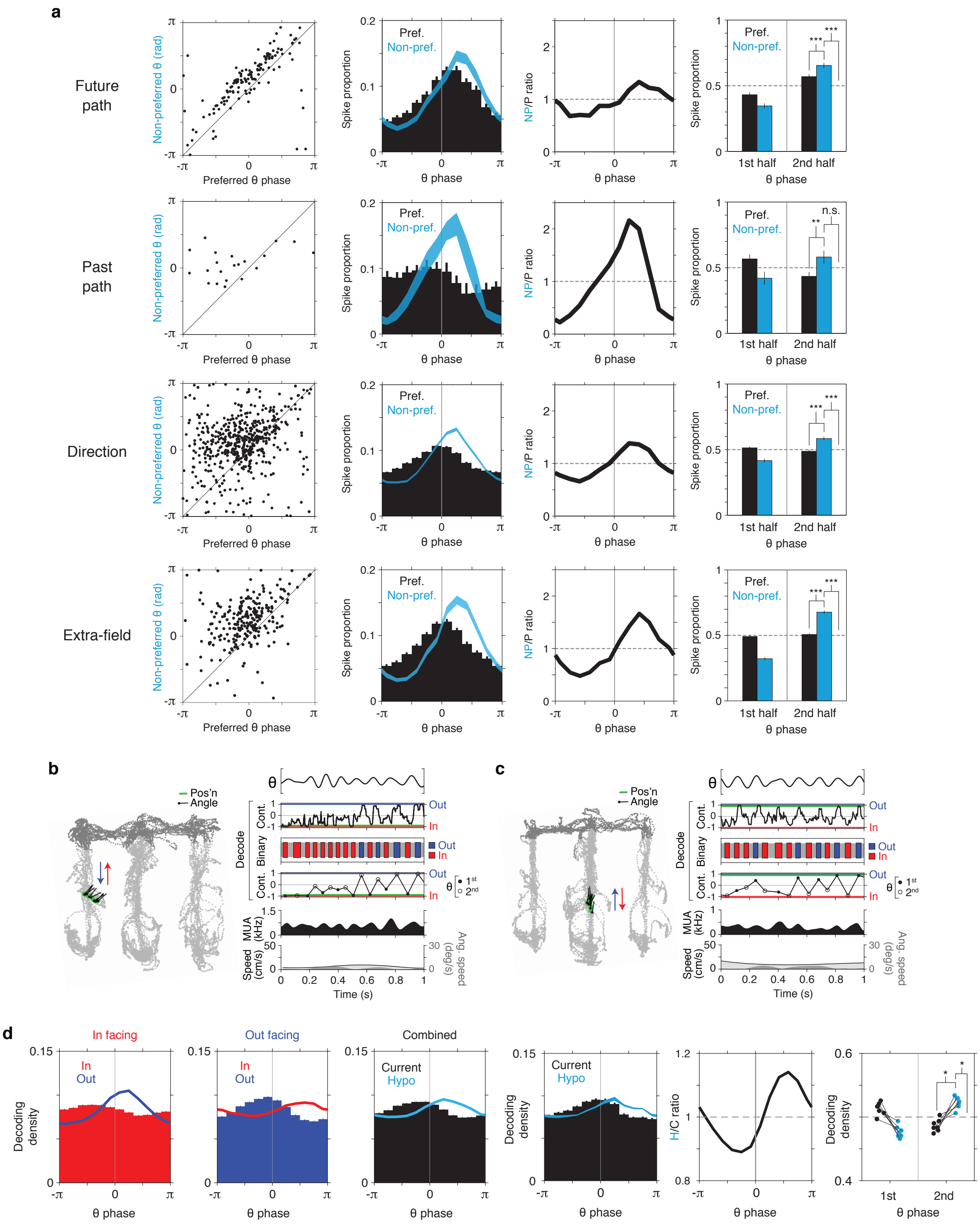

### Fig. S10. Cycling of direction at the population level.

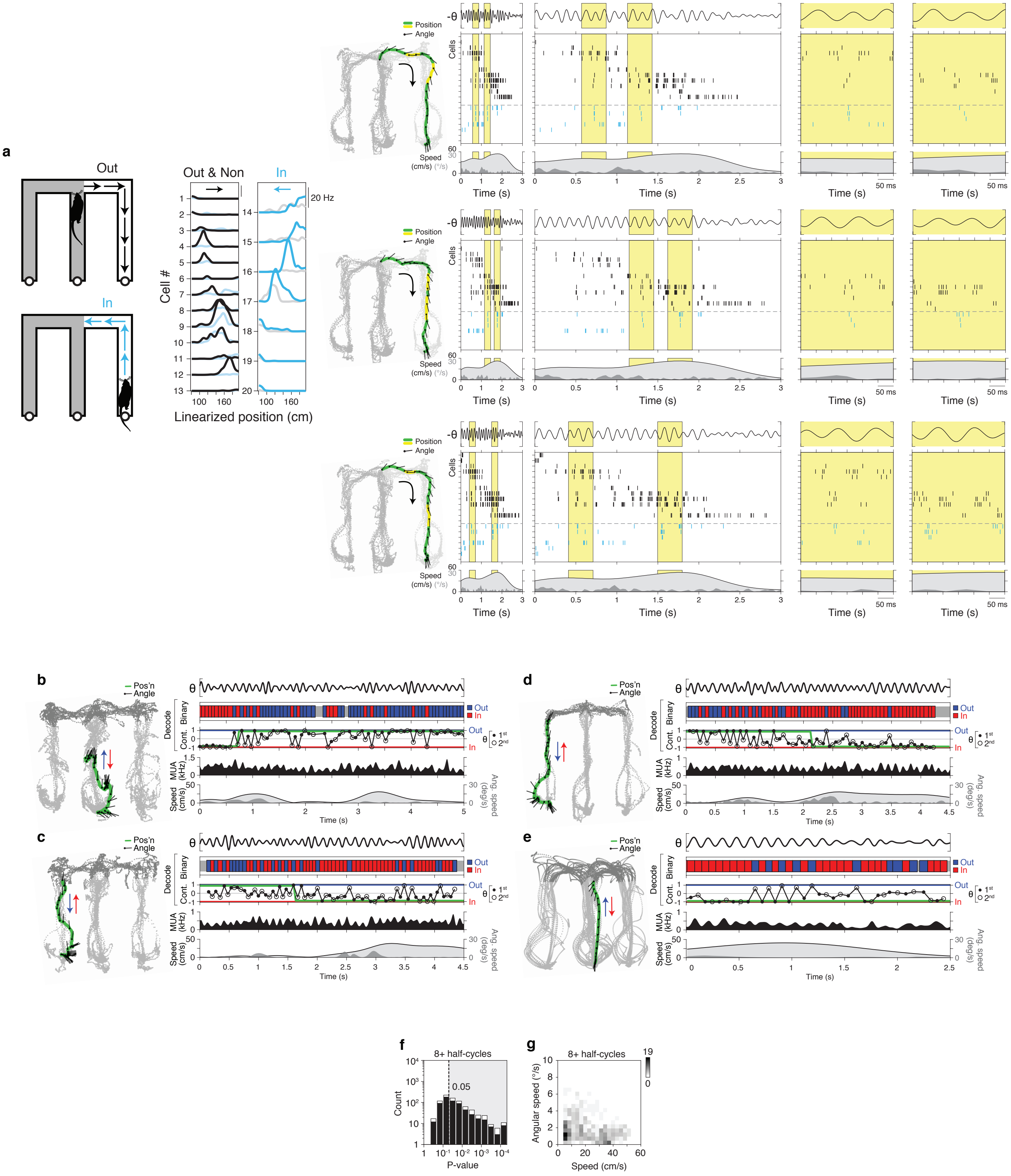
